# Maintenance of Complex I and respiratory super-complexes by NDUF-11 is essential for respiratory function, mitochondrial structure and health in *C. elegans*

**DOI:** 10.1101/2021.01.06.425530

**Authors:** Amber Knapp-Wilson, Gonçalo C. Pereira, Emma Buzzard, Andrew Richardson, Robin A. Corey, Chris Neal, Paul Verkade, Andrew P. Halestrap, Vicki A.M. Gold, Patricia Kuwabara, Ian Collinson

## Abstract

Mitochondrial super-complexes form around a conserved core of monomeric complex I and dimeric complex III; wherein subunit NDUFA11, of the former, is conspicuously situated at the interface. We identified *B0491*.*5* (*NDUF-11*) as the *C. elegans* homologue, of which animals homozygous for a CRISPR-Cas9 generated knockout allele arrested at the L2 development stage. Reducing expression by RNAi allowed development to the adult stage, enabling characterisation of the consequences: destabilisation of complex I and its super-complexes, and perturbation of respiratory function. The loss of NADH-dehydrogenase activity is compensated by enhanced complex II activity, resulting in excessive detrimental ROS production. Meanwhile, electron cryo-tomography highlight aberrant cristae morphology and widening of the inter-membrane space and cristae junctions. The requirement of NDUF-11 for balanced respiration, mitochondrial morphology and development highlights the importance of complex I/ super-complex maintenance. Their perturbation by this, or other means, is likely to be the cause of metabolic stress and disease.

## INTRODUCTION

Mitochondria are dynamic organelles operating as hubs for energy production, metabolic and biosynthetic pathways and Ca^2+^ homeostasis. Consequently, they are intrinsically associated with oxidative stress, proteostasis and cell signalling responsible for maintaining cellular and organismal health, and for survival [1-4]. They generate ATP by oxidative phosphorylation (OXPHOS) to power cellular activities [5, 6]. This is achieved by four large multimeric electron transfer complexes (CI, II, III and IV) and the ATP synthase of the inner mitochondrial membrane (IMM). The complexes I-IV form a continuous electron transfer chain (ETC), orchestrating the step-wise transfer of electrons from NADH (Complex I – NADH:ubiquinone oxidoreductase) and FADH_2_ (Complex II – succinate:ubiquinone oxidoreductase), *via* Complex III (cytochrome *bc*_*1*_) to molecular oxygen at Complex IV (cytochrome *c* oxidase). This flow of electrons – down an electro-chemical redox potential – liberates energy which is conserved by proton pumping across the IMM from the matrix to the intermembrane space. This electro-chemical gradient of protons – the proton-motive-force – is used to power the ATP synthase.

The respiratory complexes can form functional higher order assemblies with defined stoichiometries ([7] and references therein). These super-complexes, also known as the ‘respirasome’, were first observed by non-denaturing blue-native polyacrylamide gel electrophoresis (BN-PAGE) [8]. The mammalian super-complexes, visualised by cryo-electron microscopy (cryo-EM), consist of monomeric CI associated with dimeric CIII and a single CIV (SCI_1_:III_2_:IV_1_; Fig. 1 A) [9-11]. The acquisition of such a super-complex structure has lent credibility to the existence of discrete entities for the containment of the entire ETC; in contrast to the classical view of discontinuous electron transfer complexes connected by mobile electron carriers. Various super-complexes, consisting of a conserved core of SCI_1_:III_2_, have been visualised by cryo-electron tomography (cryo-ET) *in situ –* in the flat surface of cristae membranes of yeast, plants and mammals – with slight variations in the position and stoichiometry of CIV [7, 12]. This common architecture of SCI_1_:III_2_ is highly suggestive of an important functional and/ or structural role of the super-complexes. However, despite the evidence supporting the existence of structurally conserved respiratory super-complexes, the significance of these assemblies is an open question.

**Figure 1.**
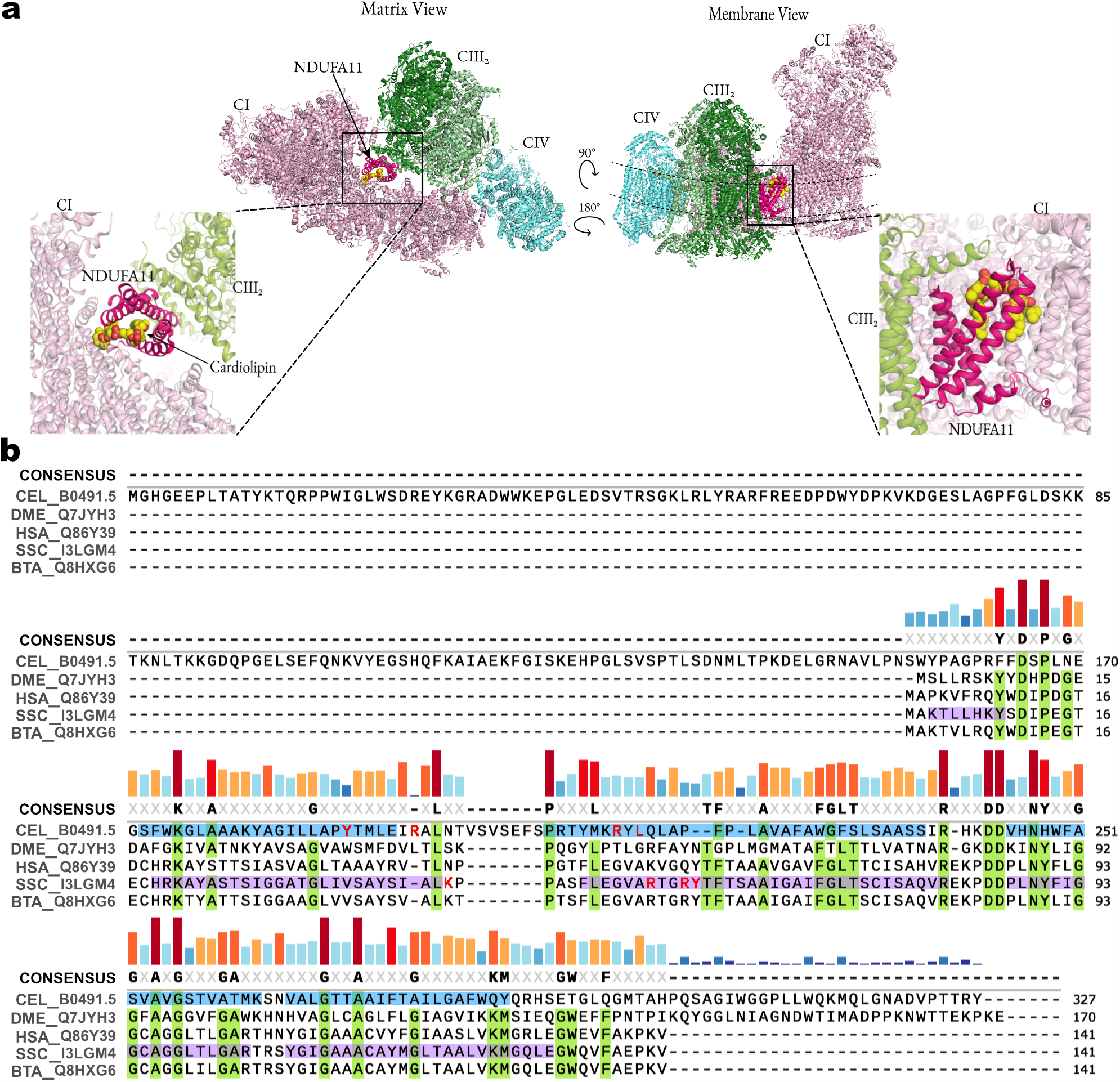
Mitochondrial complexes and supercomplexes with subunit NDUFA11. **(a)** Cryo-EM structure of the human respirasome SCI:III_2_:IV (PDB 5XTH) [80] viewed from the matrix (top left) and membrane (top right) rotated about the membrane axis. The boxed area highlights NDUFA11 and cardiolipin. CI, light pink; CIII_2_, green; CIV, blue; NDUFA11, magenta; cardiolipin, yellow. **(b)** Multiple sequence alignment of CEL-B0491.5/NDUFA11 generated by Clustal Omega [81] and graphically handled in SnapeGene. Amino acids highlighted in lilac represent TMH regions based on porcine NDUFA11 (pdb: 5GUP, [19]); while those in cyan are predicted CEL-B0491.5 TMH regions determined by JPred4 [82]. Conserved residues across entries are highlighted in light green. The sequence conservation is shown by the coloured vertical bars and a threshold of > 75% was used for the consensus. Residues important for cardiolipin binding are highlighted red. Abbreviations: CI – Complex I, CIII_2_ – dimer of Complex II, CIV – Complex IV, CL – cardiolipin, IMS – intermembrane space, THM – transmembrane helix, CEL – *Caenorhabditis elegans*, DME – *Drosophila melanogaster*, HSA – *Human sapiens*, SSC – *Sus scrofa*, BTA – *Bos taurus*.

We set out to address this problem by exploiting the genetically tractable multi-cellular model organism *C. elegans*, wherein we identified and examined the role of the homologue of the mammalian NDUFA11 – a supernumerary subunit of Complex I. NDUFA11 is an integral membrane protein positioned at the interface between Complex I and Complex III posited to support a critical interaction within the respiratory super-complex (Fig. 1A) [9-11]. However, it should be noted that NDUFA11 represents only one of these interaction points, specifically at the IMM [13]. NDUFA11 has previously been implicated in Complex I assembly [14-16] and its perturbed expression – caused by a faulty splicing – results in fatal infantile lactic acidemia, encephalocardiomyopathy and late-onset myopathy [17, 18]. It is unclear if loss off NDUFA11 activity also results in super-complex instability, or indeed if this higher order destabilisation accounts for the documented, or indeed any other, mitochondrial disease.

Our study confirms the importance of NDUFA-11 for development to adulthood and for the assembly and maintenance of Complex I and the respiratory super-complexes. Depletion of *C. elegans* NDUFA11, led to respiratory super-complex destabilisation. This in turn enabled us to assess the importance of Complex I and super-complex integrity for mitochondrial function and morphology, and the consequences of the effects of this destabilisation for whole animal physiology and health.

## RESULTS

### Identification of the *C. elegans NDUFA11* homologue

Mammalian NDUFA11 is a supernumerary subunit of Complex I – situated in the membrane arm and in contact with Complex III within the respiratory super-complex (Fig. 1A and Supp. Fig. S1a). Interestingly, it is also occupied with a tightly bound phospholipid cardiolipin (CL), situated on the matrix surface of the IMM (Fig. 1A and Supp. Fig. S1a-c). It is composed of a bundle of four transmembrane helices (TMHs) flanked by a short N-terminal helix and a C-terminal loop region. The *C. elegans B0491*.*5* gene was initially identified as encoding an *NDUFA11* homologue by psi-BLAST using the gene product sequence of the *Drosophila* counterpart (B14.7) as the query (E-value = 2e-06; identity=26%). A multiple sequence alignment of mammalian and *Drosophila* NDUFA11 proteins with worm B0491.5 highlights the conservation across homologues of membrane topology and CL binding residues (Fig. 1B). *C. elegans* B0491.5 also has a unique N-terminal extension (Fig. 1B).

Given the availability of several high-resolution structures of respiratory complexes containing NDUFA11 [9-11], the structural conservation of worm B0491.5 was explored by generating a homology model based on the cryo-EM structure of porcine NDUFA11 (pdb: 5GUP [19]) using Modeller [20] and molecular dynamics simulations (for details, refer to Material and Methods; Suppl. Fig. S1b). The derived structure is stable (Suppl. Fig. S1c) and shows the conservation of predicted features, including TMH regions and a pocket to support CL binding (Suppl. Fig. S1d, orange, right). Taken together, the sequence analysis and homology modelling confirm the classification of *C. elegan*s B0491.5, hereafter referred to as the NDUF-11 (encoded by *nduf-11*).

### Reduction of *nduf-11* expression causes growth arrest and lifespan extension

A *B0491*.*5*/*nduf-11*(*cr51*) deletion allele designed to abrogate protein-coding potential was obtained by CRISPR-Cas9 gene editing. Animals homozygous for *nduf-11*(*cr51*) – *i*.*e*. devoid of protein – arrested at the second larval stage (L2) of development, a lethal phenotype commonly displayed by animals carrying mutations in mitochondrial genes [21]. Because, as we have shown, *nduf-11(cr51)* homozygotes display an early larval arrest, it was not possible to obtain sufficient biomass for biochemical analysis. Instead, viable *C. elegans* with reduced *nduf-11* activity were recovered using feeding RNAi to enable the bulk isolation of mitochondria. Adults continuously fed with *nduf-11*(*RNAi*) were smaller and thinner and also produced fewer progeny than control mock RNAi treated *N2* adults (Fig. 2a; Table 1). However, similar to *nudf-11* knockout mutants, the progeny produced by these RNAi treated adults failed to develop into fecund adults and arrested at the L2 stage. Western blot quantification estimated that RNAi led to an ∼ 83% reduction in NDUF-11 protein levels (Fig. 2b).

**Table 1.** Brood count after *nduf-11* silencing.

|  | 2<sup>nd</sup> generation developmental arrest |  |  |  |  |  |
| --- | --- | --- | --- | --- | --- | --- |
|  | Total Brood Count<sup>a</sup> | L1 | L2 | L3 | L4 | Adult |
| Wild type (n=6) | 282 ± 2.70 (n=6) | - | - | - | - | 100 ± 0% |
| *nduf-11* (RNAi) | 1.9 ± 0.62 *** (n=10) | 12.4 ± 6.05% | 87.6 ± 6.05% | - | - | - |

<sup>a</sup> Mean ± SEM, 6 (control) and 18 (*nduf-11* (RNAi)) independent experiments, for the Developmental Stage analysis. Differences between group on the Brood Count analysis were assessed by an Unpaired t-Test. \*\*\*, $p < 0.001$ .

**Figure 2.**
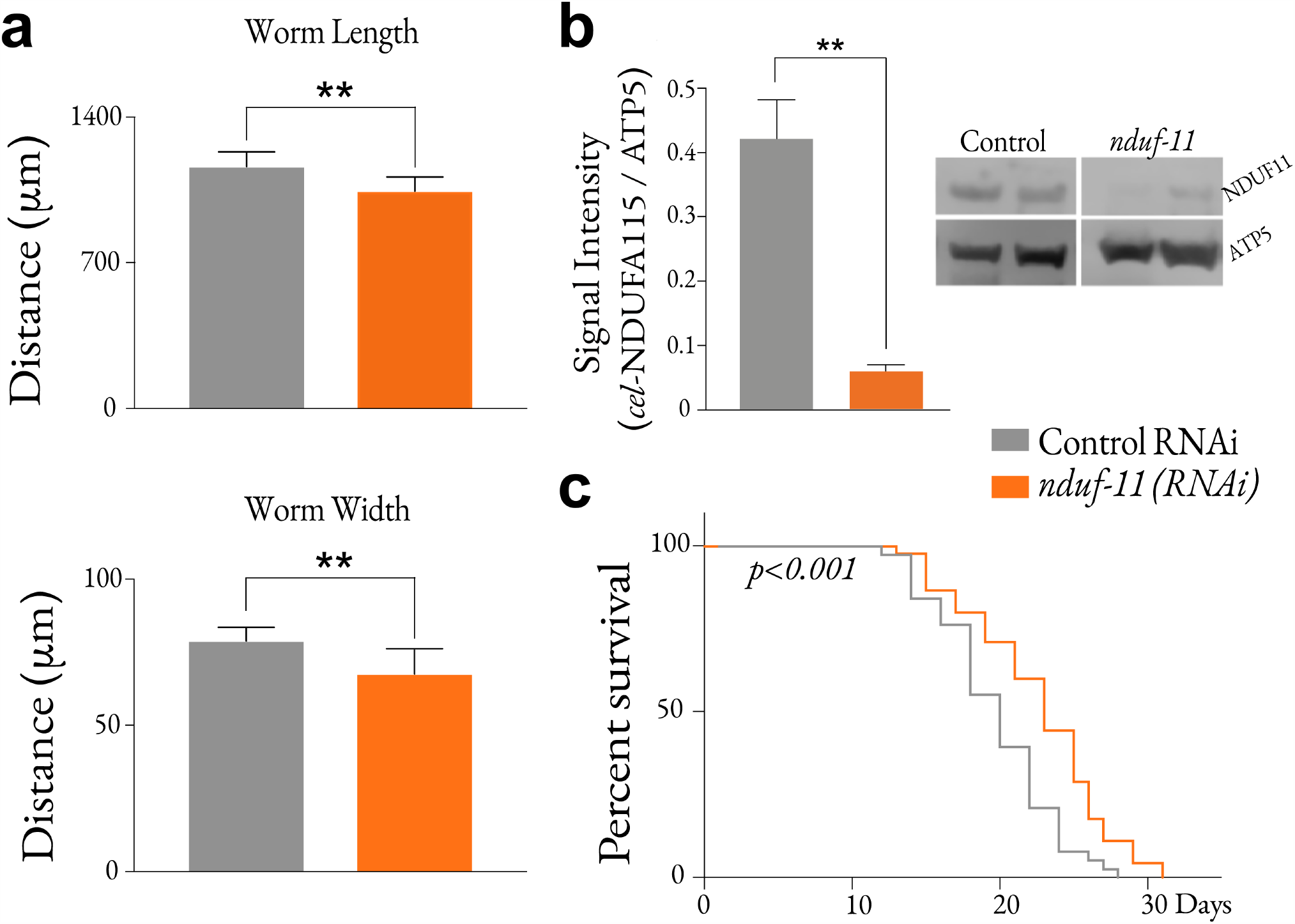
Characterisation of the *nduf-11*(RNAi). **(a)** Morphometric analysis of *nduf-11*(*RNAi*) adult worms. n= 11 (wild type) and n=8 *nduf-11*(*RNAi*) adults were analysed. Data shown as mean±SEM. Statistical analysis, Students’ t-test. **, p < 0.01. **(b)** Reduction of NDUF-11 protein after RNAi treatment in mitochondria. 50 µg of isolated mitochondria were loaded per lane. Data shown as mean±SEM of n=5. Statistical analysis, Students’ t-test. **, p < 0.01. **(c)** *nduf-11(RNAi*) is responsible for a lifespan extension. Wild-type (n=45); *nduf-11*(*RNAi*) (n=38). Kaplan-Meier survival statistics.

Continuous application of *nduf-11*(*RNAi*) also led to an extension in lifespan when compared to mock RNAi treated control animals (Fig. 2c), confirming a previous observation [22]. These analyses confirm that *nduf-11 (RNAi*) treatment is effective in reducing the corresponding protein levels, while allowing worms to develop into fecund adults.

### Analysis of the mitochondrial and cytosolic proteome following NDUF-11 depletion by RNAi

The underlying cause of the lethality of the *nduf-11* knockout was assessed through an examination of the less severe phenotype observed by reducing *nduf-11* expression by RNAi treatment. This analysis began by an investigation of the corresponding changes in the mitochondrial and cytosolic proteomes. For this purpose, mitochondrial and cytosolic fractions were collected from whole worms (see Methods for details); the latter being, as much as possible, devoid of nuclear, plasma membrane and other endosomal systems.

Quantitative mass spectrometry comparing native and depleted levels of NDUF-11 protein identified 6440 proteins in each fraction of which less than 10% were significantly up- or down-regulated (Fig. 3a). Of the affected proteins, the majority were downregulated upon NDUF-11 depletion: 4.85% and 3.26% attaining statistical significance for the mitochondrial and cytosolic fractions, respectively (Fig. 3b). In contrast, only 2.87% and 1.24% were upregulated in the corresponding mitochondrial and cytosolic fractions (Fig. 3b). As expected, the data confirmed NDUF-11 was downregulated to about 20 % of wild-type levels (Fig. 3c, blue dot) – similar to the value estimated by Western blotting (Fig. 2b). The remaining Complex I subunits were downregulated by about 50% (Fig. S2), suggesting that NDUF-11 is indeed important for Complex I integrity, as previously documented [14]. Conversely, other respiratory complex subunits were marginally upregulated, albeit not above a statistically significant threshold (Fig. S2).

**Figure 3.**
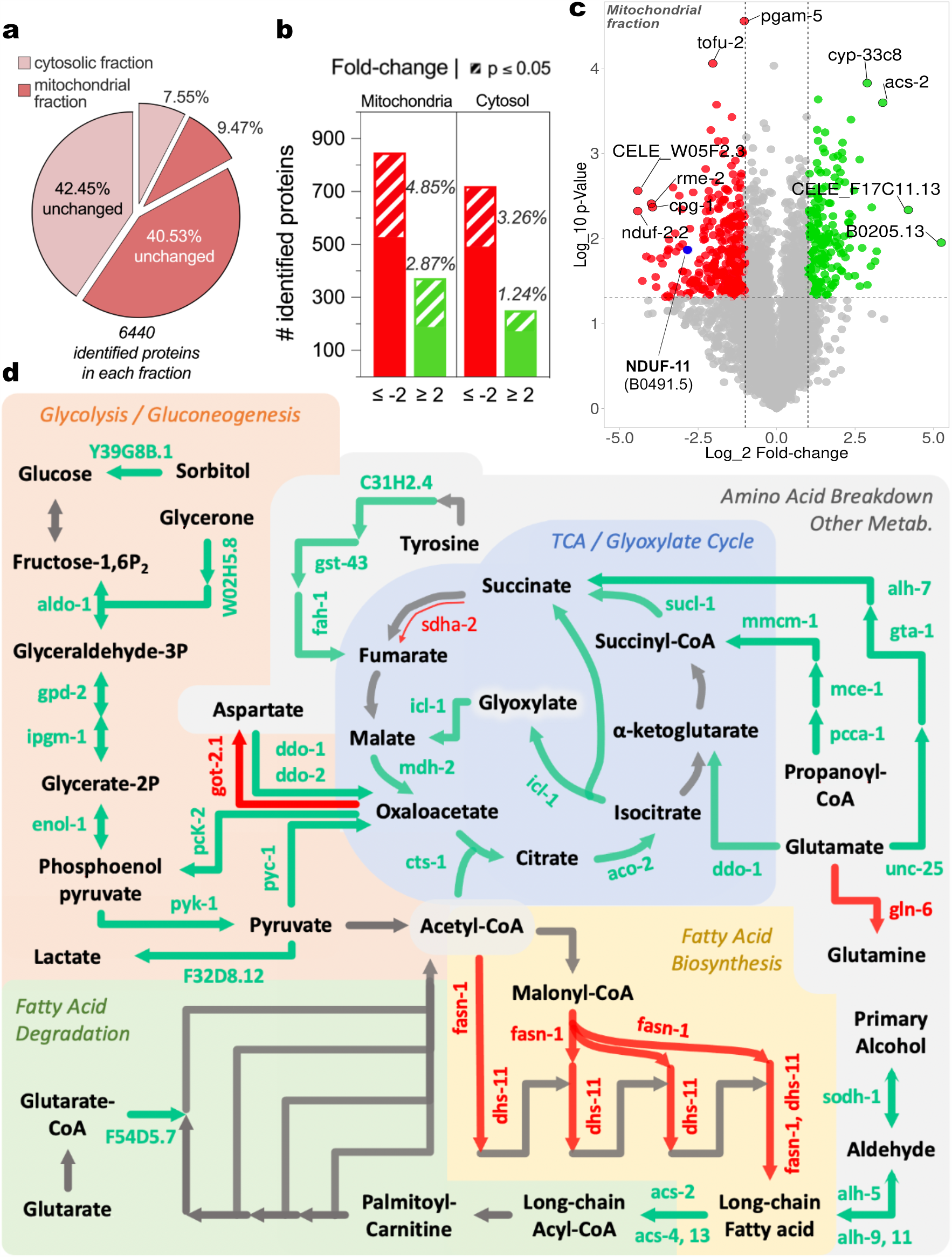
**(a-b)** Summary data of the mass spectrometry analysis of the cytosolic and isolated mitochondrial fractions from control and *nduf-11*(RNAi) groups. The exploding slices on the pie-chart displayed on panel **a** represent a subset of proteins which were up/downregulated. The bar chart on panel **b** sorts these subsets of proteins accordingly to their levels (up or downregulated, green or red colour, respectively) and p-value. n=3. **(c)** Volcano-plot showing protein expression changes in isolated mitochondria after RNAi treatment. NDUF-11 is highlighted in blue on the plot. Protein significantly upregulated are shown in green while those downregulated are shown in red. Protein which levels are similar to control levels are depicted in grey. The top 10 most significant hits are also labelled. n=3. **(d)** Summary of the major changes in several metabolic pathways after RNAi treatment. The significantly up- and downregulated proteins from the cytosolic and mitochondrial fractions shown in panel a-b were submitted through a KEGG analysis. The majority of changes were observed in metabolic pathways which are depicted here in boxes with different background (glycolysis/gluconeogenesis, amino acid breakdown, fatty acid biosynthesis and breakdown, tricarboxylic acid cycle and glyoxylate cycle). Green arrows represent proteins which were upregulated upon *nduf-11 (RNAi)* silencing and, conversely, red arrows represent downregulated proteins. Grey arrows represent metabolic steps in the depicted pathway but which protein levels remain unchanged. Thin arrow represents germline-specific isoforms.

In addition to changes in the levels of the ETC constituents, we observed downregulation of proteins responsible for fatty-acid biosynthesis, and an upregulation in the rate-limiting enzyme of β-oxidation – acyl-CoA synthetase (ACS; encoded by acs-2 in *C. elegans*); indicative of an upregulation of the fatty acid catabolic pathway (Fig. 3d). Furthermore, the data suggests a strong remodelling of the TCA cycle towards a glyoxylate cycle. In this regard, amino acid breakdown and the propanoyl-CoA pathway is upregulated to replenish intermediates in the TCA/ glyoxylate cycle. Finally, several enzymes of the glycolysis/ gluconeogenesis axis are upregulated after NDUF-11 depletion. It is noticeable that both *pyk-1*, rate controlling in glycolysis, and the *pyc-1/ pck-2* pair, rate controlling in gluconeogenesis, are upregulated, suggesting a possible ‘futile metabolic cycle’. However, considering that our sample is from a pool of cells from the whole organism, it is reasonable to assume that different tissues might respond differently to reduced levels of NDUF-11. In this regard, we observed a downregulation of *sdha-2* (Fig. 3d and Fig. S2) and *asb-1* (Fig. S2), which are germline-specific isoforms of Complex II and Complex V, respectively.

### Confirmation that NDUF-11 is a constituent of *C. elegans* mitochondrial Complex I

Experiments were conducted to confirm that the *C. elegans* NDUFA11 homologue is associated with Complex I. Mitochondria from animals with native and depleted levels of NDUF-11, were analysed by BN-PAGE, using conditions optimised for detecting respiratory complexes and ATP synthase. As a first step, the membranes were solubilised harshly with the detergent Triton X-100 (TX-100) prior to non-denaturing BN-PAGE (Fig. 4a, left). The protein complexes were visualised in this first dimension by Coomassie staining (Fig. 4a, left), and Complex I by an in-gel activity assay [23] (Fig. 4a, right). In the former, the two most prominent bands were monomeric forms of Complex I and the ATP synthase, which had been separated from their respective respiratory super-complex and homo-dimeric forms by TX-100. Individual proteins of complexes resolved by BN-PAGE were separated in a 2^nd^ dimension by denaturation with LDS-PAGE. The proteins were then visualised by silver staining and immuno-blotting, identifying NDUF-11 as a component of the monomeric Complex I (Fig. 4b). As predicted, NDUF-11 was reduced in samples obtained from *nduf-11* RNAi-treated mitochondria. This analysis also showed there is a corresponding lower recovery of intact and active Complex I from extracts where NDUF-11 is depleted, while the ATP synthase is unaffected, suggesting that monomeric Complex I without NDUF-11 is unstable when solubilised by TX-100.

**Figure 4.**
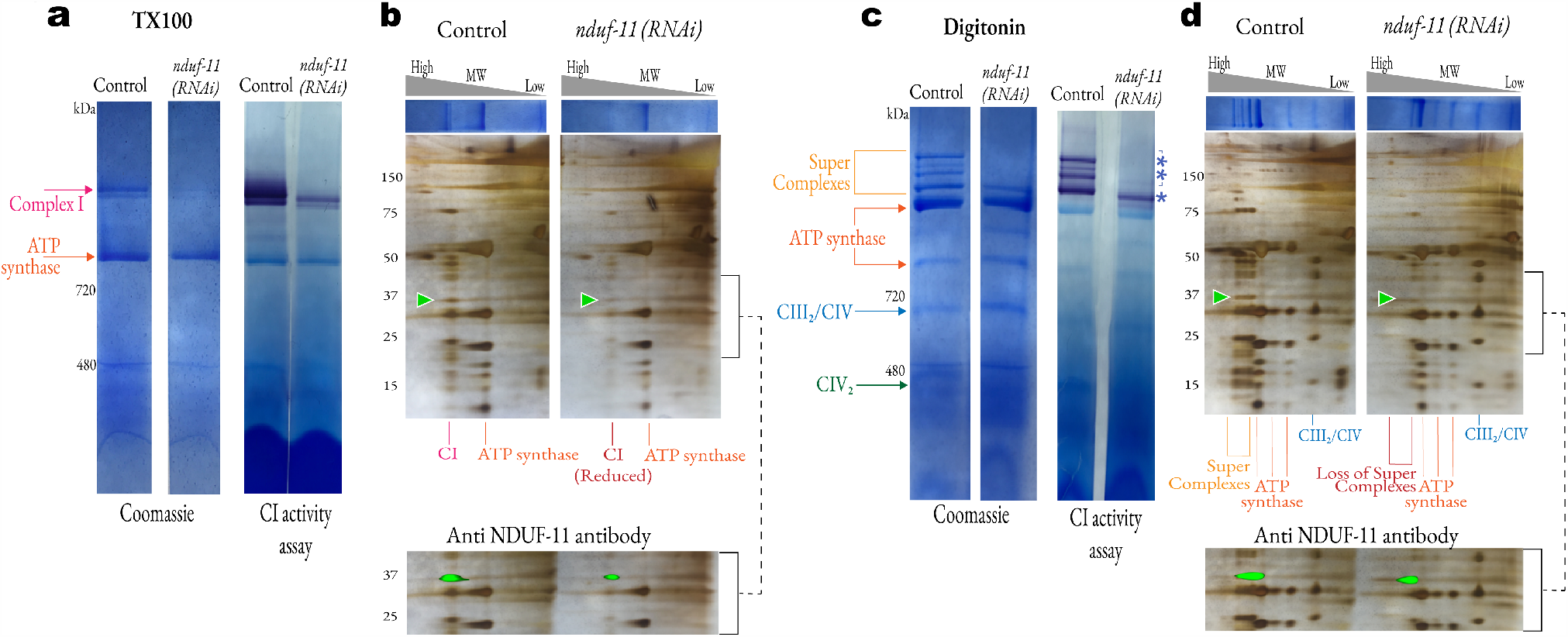
Analysis of mitochondrial respiratory complexes. **(a)** BN-PAGE analysis of TX-100 extracts of mitochondrial proteins. 78 µg of solubilised mitochondria were loaded per lane. Top arrow, monomeric CI; bottom arrow, ATP synthase (CV) **(b)** 2D BN-PAGE analysis of TX-100 mitochondrial protein extracts. 1^st^ dimension BN-PAGE Coomassie stained gel lane (high-low MWt, left to right) resolved on denaturing LDS-PAGE gel and silver stained. Green arrowhead indicates the position of the NDUF-11. Inset, antibody detection of NDUF-11 superimposed on silver-stained gel. The signal from the protein ladder in both images was used to align the superimposed result. **(c)** BN-PAGE analysis of digitonin extracts of mitochondrial proteins. 150 µg of solubilised mitochondria were loaded per lane. Positions of respiratory complexes were determined by in-gel activity assays (for CI, IV and V; data not shown). Double asterisk represents the loss of higher molecular weight super-complexes in the *nduf-11(RNAi)* group. Single asterisk represents the only detectable form of super-complex in the *ndufa-11(RNAi)* group, probably representing the SCI_1_:CIII^2^ assembly. **(d)** 2D BN-PAGE analysis of digitonin mitochondrial protein extracts. 1^st^ dimension BN-PAGE Coomassie stained gel lane (high-low MWt, left to right) resolved on denaturing LDS-PAGE gel and silver stained. Green arrowhead indicates the position of the NUDF-11. Inset, antibody detection of NDUF-11 superimposed on silver-stained gel. The signal from the protein ladder in both images was used to align the superimposed result.

### Reduction of NDUF-11 leads to destabilisation of the respiratory super-complexes

The integrity of the respiratory super-complexes during BN-PAGE is partially dependent on the nature of the detergent used for solubilisation [23]. Thus, for these studies, TX-100 was replaced by the milder detergent digitonin, known to preserve the super-complexes containing Complex I [24-26] (Fig. 4c). Equivalent proportions of extracts derived from mitochondria with wild-type and depleted levels of NDUF-11 were analysed by BN-PAGE. A series of high molecular weight respiratory super-complexes were visualised by a combination of Coomassie blue and in-gel Complex I activity staining in the native extract (Fig. 4c). Conversely, NDUF-11-depleted samples contain only the lower molecular weight form of Complex I, and critically are devoid of higher molecular weight super-complexes seen in the control (Fig. 4c; lower asterisk *versus* upper two asterisks). Both protein (Coomassie) and Complex I activity staining show that the total amount of Complex I is reduced to levels consistent with the proteomic analysis (Fig. S2). The quantity and state of the ATP synthase were unaffected (Fig. 4c).

Second-dimension denaturing LDS-PAGE confirms the presence of Complex I subunits in the different bands assigned to high-molecular weight respiratory super-complexes (Fig. 4d). Immunoblotting confirmed the presence of NDUF-11 in all of these super-complexes (Fig. 4d, lower panel).

Taken together, the results suggest that depletion of NDUF-11 destabilises respiratory super-complexes resulting in either their fragmentation within the membrane or their loss under mild extraction conditions.

### Reduction of NDUF-11 affects mitochondrial morphology

A targeted GFP reporter was used to observe changes in mitochondrial morphology and distribution in body wall muscles. Compared to wild-type animals, NDUF-11 depleted mitochondria appeared fragmented and less reticulated (Fig. 5a); a possible indicator of a pathological condition. These morphological differences were also evident after inspection using (Fig. 5b) transmission electron microscopy (TEM) of high-pressure frozen and freeze substituted *C. elegans* worms. The analysis showed that NDUF-11 depleted mitochondria were more spherical compared to the control, which had a more elongated shape (Fig. 5b, right panels).

**Figure 5.**
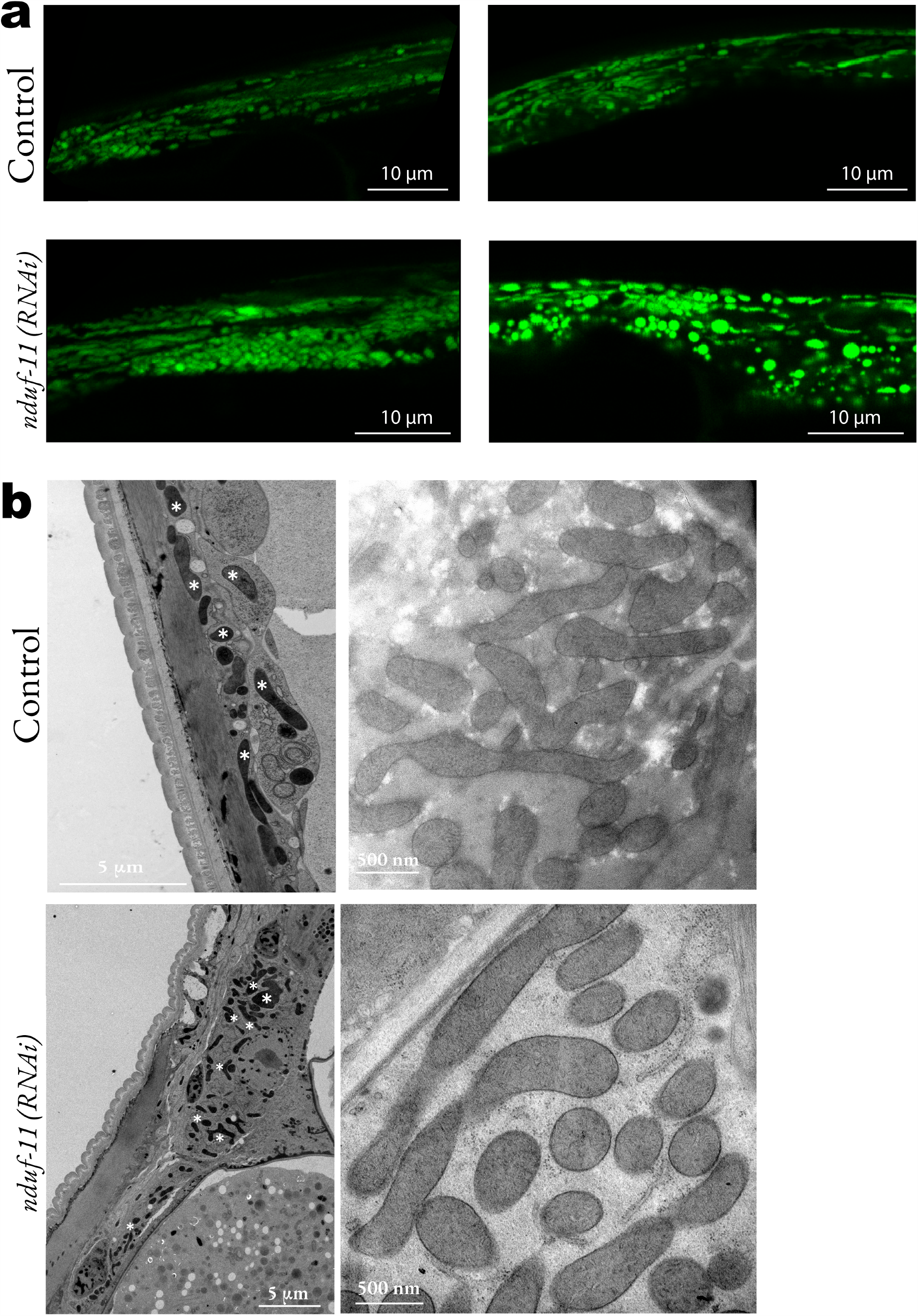
Mitochondrial morphology of *nduf-11(RNAi*). (**a**) Representative images of confocal microscopy of GFP-labelled mitochondria in the body wall muscles (Pmyo-3::mito::GFP) of wild type and *nduf-11*(*RNAi*) animals. A total of 20 worms were imaged in three independent experiments. **(b)** Electron micrographs of body wall mitochondria in wild type and *nduf-11*(*RNAi*) adult animals. Images are representative of 7 worms in each group of a single experiment. Some but not all mitochondria in the micrograph have been highlighted by white asterisks on the left hand-side panel.

In order to explore the mitochondrial interior and ultrastructure in greater detail, samples of isolated mitochondria were vitrified and subjected to electron cryo-tomography (cryo-ET). The tomographic reconstructions (controls: Fig. 6a-c; Movie M1, and RNAi treated: Fig. 6d-f; Movie M2) show that mitochondria with depleted NDUF-11 – with reduced levels of Complex I and destabilised super-complexes – contain a higher number of identifiable cristae compared to control specimens (Fig. 6i). Interestingly, the frequency of cristae junctions (CJ) is similar to the control (Fig. S3a), suggesting that a high number of cristae are fragmented and not connected to the IMM. This can be seen directly in the tomographic images (Fig. 6d-f, marked with asterisks in panel d), where cristae from NDUF-11 depleted mitochondria lose the classical mitochondrial lamellar morphology, resulting in disconnected sac-like crista with a lower surface area to volume ratio (Fig. 6j). Consequently, the total cristae surface area and volume are increased (Fig. S3b-c) despite the unchanged average dimensions (surface area or volume) of them individually (Fig. S3d-e).

**Figure 6.**
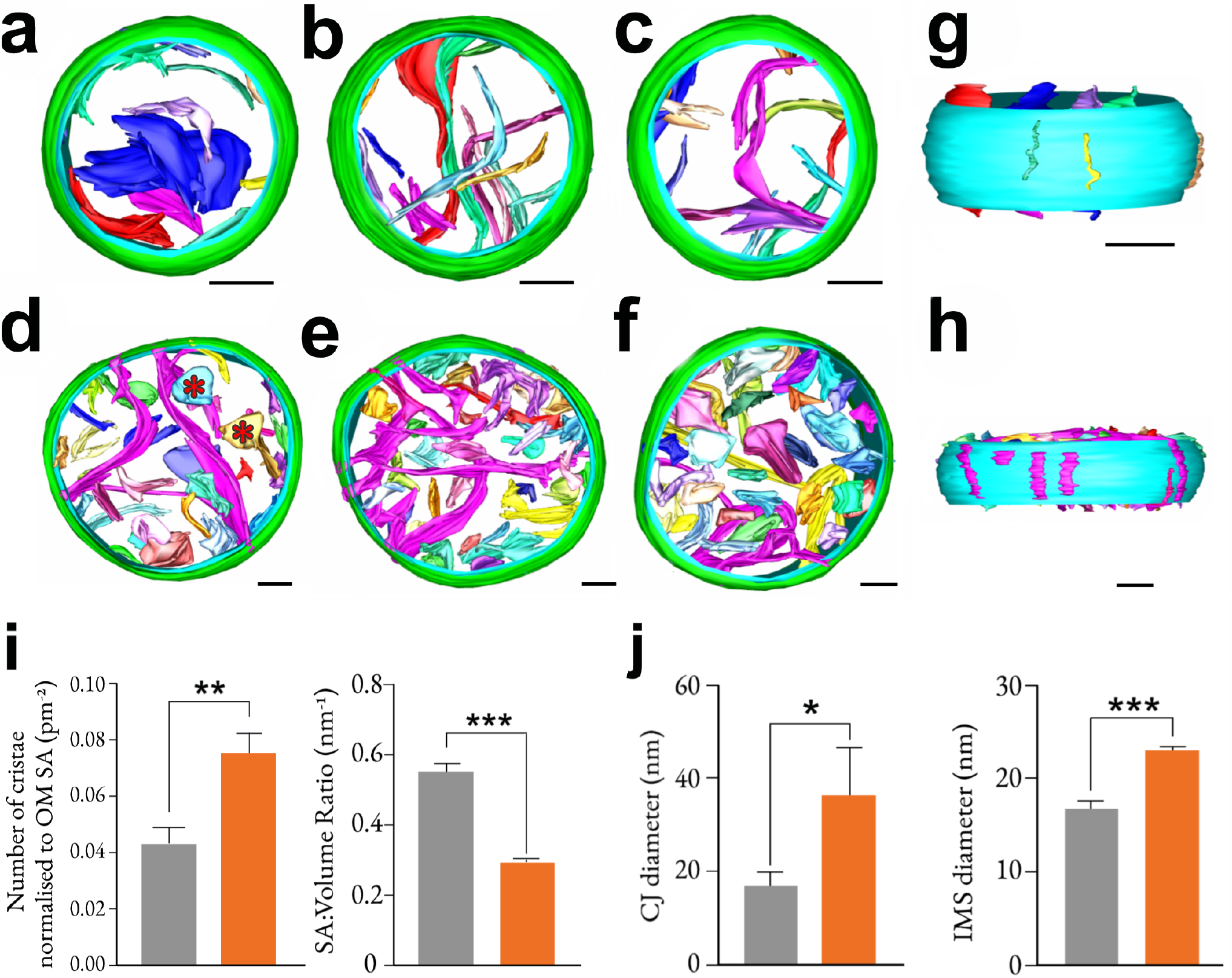
NDUF-11 knockdown affects formation of lamellar cristae. **(a-f)** Tomographic reconstruction and segmentation of representative wild-type **(a-c)** and *nduf-11*(RNAi) **(d-f)** mitochondria isolated from *C. elegans*. Wild-type mitochondria have lamellar cristae, whereas *nduf-11* (RNAi) mitochondria have more sac-like cristae, as indicated by the red asterisks in (d). Green outer-membrane, cyan inner-membrane, each crista membrane has a different colour. Reconstructions were made in IMOD 4.9. Scale bars: 100nm. See supplementary Movie M1 (control) and M2 (RNAi). **(g-h)** Side view of a wild-type and *nduf-11*(RNAi) mitochondrion from (**a**) and (**e**), respectively, with outer-membrane hidden, revealing slit-like crista junctions characteristic of *C. elegans* mitochondria. Junctions in *nduf-11*(RNAi) mitochondria are visibly wider than those in the wild-type. Scale bars: 100nm. **(i)** Crista SA: Volume ratio. Mesh surface area and volume inside the mesh of each membrane from mitochondrial reconstructions shown in panels a-f were calculated computationally (n=3 mitochondria for each condition, and n = 37 (wild-type) and 105 (NDUF-11) for total crista analysed). **(j)** Number of cristae per mitochondrion (normalised to outer membrane surface area) from wild-type and *nduf-11*(RNAi) mitochondrial reconstruction shown in panels a-f (n=3 mitochondria for each condition). **(k)** Diameter of CJs. CJs were measured from the same data used to generate the images in panels a-f (n=3 mitochondria for each condition). **(l)** IMS diameter. IMS was measured from the same data used to generate the images in panels a-f (n=3 mitochondria for each condition, with 4 points per mitochondrion analysed). Data were analysed by an unpaired, parametric Student’s t test. * p ≤ 0.05, ** p ≤ 0.01, *** p ≤ 0.001. Error bars: SD.

Finally, perturbation of Complex I and its super-complexes correlated with a widening of the CJ (Fig. 6g, h, k and Fig. S3f), and an increased separation between the outer and inner membranes, widening the IMS (Fig. 6l). Since the MICOS complex, located at crista junctions, is known to mediate cristae and IMM remodelling [27] we revisited the mass spectrometry dataset (Fig. 3a-c) to assess the expression levels of key components [28], namely, IMMT-1, IMMT-2, MOMA-1, CHCH-3, F54A3.5 and W04C9.2. Only IMMT-2, a homologue of Mic60, was significantly downregulated after RNAi treatment (−2.15 log_2_ fold-change, p = 0.003 in mitochondrial fraction). Though, it should be noted that the *C. elegans* Opa1 homologue EAT-3 remained unchanged after *nduf-11* silencing.

Overall, NDUF-11 and its loss induce profound changes in the internal membrane architecture of mitochondria, possibly as a consequence of metabolic remodelling.

### Maintenance of Complex I and its super-complexes is required for optimised mitochondrial respiration and prevention of excessive ROS production

Next, we explored the consequences of NDUF-11 depletion, and subsequent respiratory complex destabilisation, on bioenergetic fitness. Thus, the respiratory performance of isolated mitochondria was assessed on the basis of their oxygen consumption and membrane potential. Upon addition of the exogenous Complex I substrates pyruvate/ malate, similar rates of basal respiration were measured for mitochondria with native or depleted levels of NDUF-11 (Fig. 7a, top panel). When stimulated by the addition of ADP, the rate of respiration increased to drive oxidative phosphorylation. This activation dropped from 6.9-fold to a modest 2.3-fold in mitochondria with reduced NDUF-11 levels. In both cases, the addition of exogenous cytochrome *c* during ADP-sustained respiration (Fig. 7 a, bottom panel) elicited little change, indicating the outer-membrane is intact and had retained endogenous cytochrome *c*. Therefore, the lowered respiratory capacity is not a consequence of mitochondria outer membrane permeabilisation. In contrast to mitochondria with wild-type levels of NDUF-11, those depleted of NDUF-11 also failed to respond to the uncoupler CCCP after addition of oligomycin. This suggests that the impairment was associated with the ETC and not the phosphorylation system.

**Figure 7.**
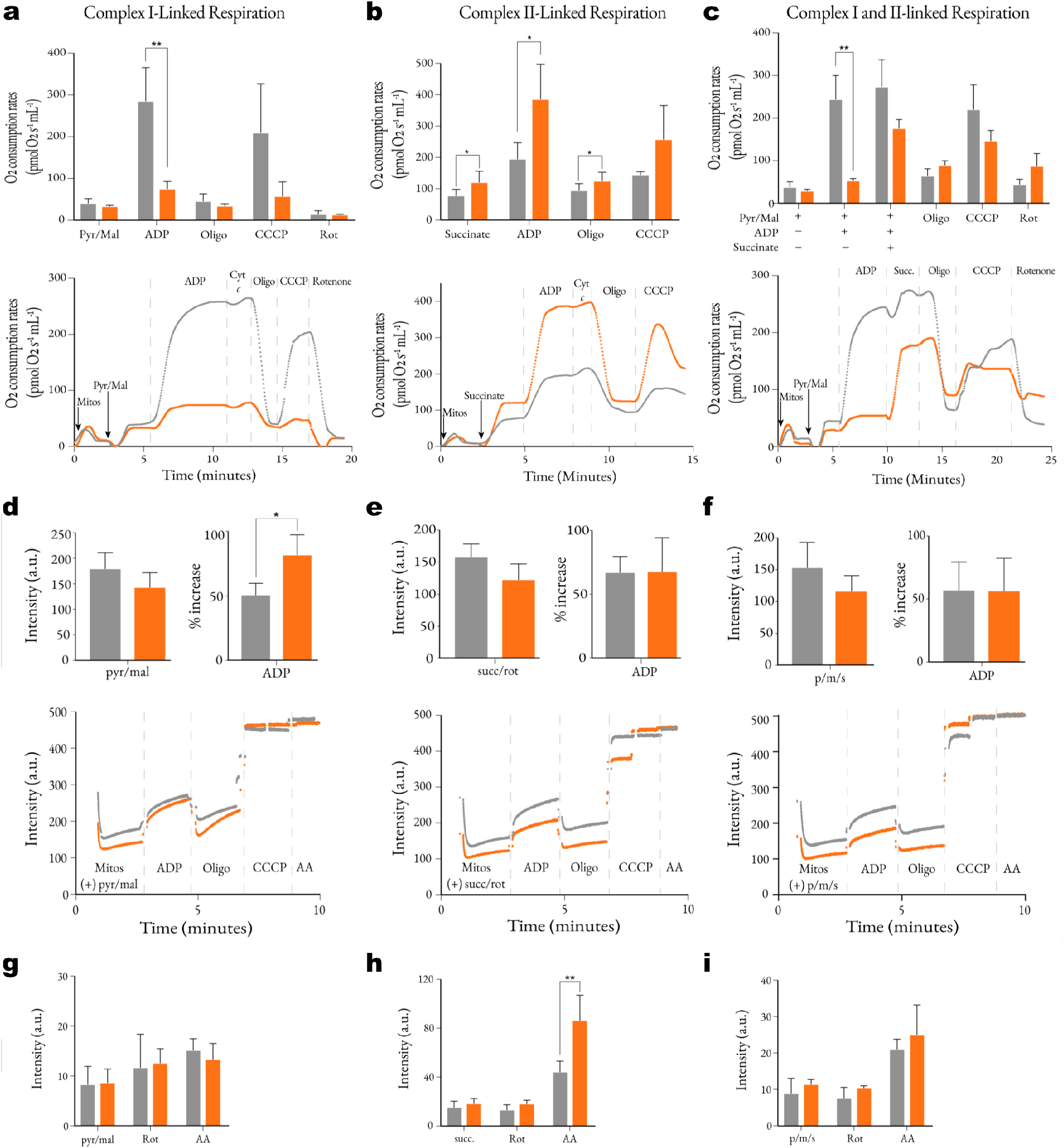
Bioenergetic analysis of mitochondria after *nduf-11(RNAi*) treatment. (**a-c**) Assessment of mitochondrial respiration by monitoring oxygen consumption in isolated fractions. Mitochondria were energised with either Complex I-linked substrates (pyruvate/malate; a), Complex II-linked substrates (succinate; b) or both (c); and, ADP, inhibitors and uncoupler were sequentially added to assess mitochondria fitness (dashed lines). Data on the line plot shows averaged values of 3-5 independent experiments. Data on bar charts is shown as mean ± SEM of 4 independent experiments. Differences between groups for each respiratory state was assessed by a paired Student’s t-test. (**d-f**) Assessment of mitochondrial respiration by monitoring membrane potential with TMRM in isolated fractions. Mitochondria were energised as in (a-c). Data on the line plot shows averaged values of TMRM signal from 3-5 independent experiments. Data on bar charts is shown as mean ± SEM of 4 independent experiments. Differences between groups for each respiratory state was assessed by a paired Student’s t-test. (**g-i**) Production of hydrogen peroxide production by isolated mitochondria monitored using Amplex Red. Mitochondria were energised as in (a-c). Data on bar charts is shown as mean ± SEM of 4 independent experiments. Differences between groups in the presence of each inhibitor was assessed by a paired Student’s t-test. *, p < 0.05; **, p < 0.01. Abbreviations: AA – antimycin A, Cyt c – cytochrome c, Pyr - pyruvate, Mal – malate, Mitos – mitochondria, Oligo – oligomycin, Rot - rotenone.

To confirm that the effects described above were specific to Complex I, its activity was inhibited by rotenone and mitochondria were energised with the Complex II substrate succinate in a separate experiment (Fig. 7b). Under these conditions, NDUF-11 depleted mitochondria responded to the addition of both ADP and CCCP – by bypassing the compromised Complex I. Surprisingly, the Complex II rescued respiration rates were consistently higher when compared to the control (Fig. 7b). In this case, the respiratory control ratio (RCR; ADP-stimulated over oligomycin-inhibited respiration), an indicator of the coupling efficiency between respiration and ADP phosphorylation, was significantly higher when NDUF-11 was depleted (3.06 ± 0.12 *versus* 2.07±0.15, p=0.002).

The data show that mitochondria with depleted NDUF-11 levels are respiration competent, and suggest that increased succinate levels could rescue mitochondrial respiration *in vivo*. To test this notion, mitochondria were incubated in pyruvate/ malate and ADP prior to the addition of succinate in the absence of rotenone to better represent those conditions experienced by mitochondria *in vivo* (Fig. 7c). Succinate addition produced only a minor rate increase in unaffected mitochondria, but was able to restore ADP-sustained respiration rates in NDUF-11 – Complex I compromised mitochondria – albeit not to wild-type values. Therefore, it can be speculated that tissues able to modulate their metabolism towards succinate production, as globally shown by our proteomic data (Fig. 3d), are better able to compensate for the loss of Complex I/ super-complexed upon *nduf-11* silencing.

To continue, the trans-membrane membrane potential of the inner membrane (ΔΨ) was investigated using the fluorescent potentiometric probe TMRM (Fig. 7d-f). The maximum observed levels of ΔΨ were approximately the same, irrespective of levels of NDUF-11 or choice of the respiratory substrate (Complex I- or Complex II-linked). However, depolarisation by the addition of ADP was greater in NDUF-11 depleted mitochondria compared to the control when mitochondria were energised with Complex I-linked substrates (Fig. 7d). ΔΨ_ADP_ reflects the equilibrium attained between PMF utilisation by the ATP production machinery (proton influx) and its generation by the ETC (proton pumping). Therefore, the reduced ability of mitochondria depleted in NDUF-11 to respire through Complex I leads to a greater reduction in ΔΨ upon ADP addition because the ETC pumps fewer protons by comparison to those with wild-type levels. This is in contrast to Complex II-linked respiration, which in this respect is unaffected (Fig. 7e). When pyruvate, malate and succinate were simultaneously added under ‘combined’ respiration conditions (Fig. 7f), ADP depolarisation was also unaffected. The data confirm that bypassing Complex I through the addition of succinate rescues the reduced bioenergetic fitness attributed to compromised Complex I and respiratory super-complex maintenance due to limited levels of NDUF-11.

We next measured hydrogen peroxide production using Amplex Red to determine if the impaired respiration displayed by the compromised mitochondria led to an elevation in ROS by the ETC. When mitochondria were energised through Complex I by pyruvate/ malate, hydrogen peroxide production was unaffected even in the presence of the ETC inhibitors rotenone (of Complex I) and antimycin A (Complex III) (Fig. 7g). By contrast, when energy was supplied *via* Complex II with succinate the mitochondria depleted in NDUF-11 experienced a burst of hydrogen peroxide production, likely caused by the increased flow of electrons through the ETC (Fig. 7h).

## DISCUSSION

NDUFA11 is one of the many supernumerary subunits of mammalian respiratory Complex I, with homologues in *Neurospora crassa* (21.3b) [29] and plants (B14.7) [30]. Interestingly, NDUFA11 also shares homology with the central and essential Tim17 family [31], including Tim17 and Tim23 of the TIM23-complex and Tim22 of the TIM22-complex. The importance of NDUFA11 as Complex I subunit was established in *N. crassa* and human cell culture studies, which revealed that impaired NDUFA11 activity led to the incomplete assembly of Complex I [14, 15]. NDUFA11 resides between assembly modules of Complex I, where it has been suggested to somehow facilitate their assembly [14, 32-35].

NDUFA11 also sits at the interface between Complexes I and III of the respiratory super-complex [9-11], in permanent association with cardiolipin [19]. This unusual lipid, first identified in mitochondria [36], is known also to be important for the structure and activities of the ETC [37-39] and ATP synthase [40, 41]. Thus, the proximity of cardiolipin to NDUFA11 is further suggestive of a critical role on the structure and function of the respiratory super-complex.

NDUFA11 was identified in the multi-cellular model organism *C. elegans* (NDUF-11) and targeted to destabilise Complex I as well as to interfere with its ability to form respiratory super-complexes. This in turn enabled an exploration of the consequences of this disruption from a biochemical perspective through to whole organism physiology. The results highlight the importance of ETC organisation for respiratory fitness, mitochondrial ultra-structure and function, and ultimately healthy development.

Perhaps unsurprisingly, deletion of *nduf-11* caused a developmental arrest, and its depletion disrupted Complex I activity with a reduction in Complex I levels and a corresponding loss of recovery of respiratory super-complexes. These results are consistent with previous work indicating that the integrity of Complex I and the respiratory super-complexes are inter-dependent [35]. They are also in agreement with NDUFA11 being an intrinsic assembly and stabilisation factor for Complex I and the respiratory super-complexes [14, 32]. They also support studies elsewhere suggesting that loss of the interface between Complex I/ Complex III renders the complex immature and prone to degradation [42, 43].

The milder phenotype displayed by worms with a reduction, but not elimination of NDUF-11, enabled us to produce sufficient biomass for biochemical studies. This, in turn, allowed us to investigate how Complex I, when compromised in activity, structure and super-complex formation, affects physiology and health in a multi-cellular organism. Despite the reduction of Complex I activity and apparent loss in its ability to organise into super-complexes, the bioenergetic performance baseline of the affected mitochondria remained largely unchanged. The effects were only revealed when mitochondria were challenged by an increase in energy-demand. This is particularly easy to assess *in vitro* in isolated mitochondrial fractions by incubation with exogenous ADP or, alternatively, by addition of uncouplers which also leads to mitochondrial depolarisation. Overall, the observed limitation in respiratory activity could explain why worms displayed a developmental arrest phenotype – discussed below.

Succinate is able to bypass and rescue (*via* Complex II) the Complex I-specific deficiencies, indicating that other respiratory complex assemblies must persist more or less intact, consistent with the blue-native gels and proteomic data. However, when succinate was added exogenously to mitochondria isolated from *nduf-11 (RNAi)* animals, a drastic increase in respiration was observed, indicating that significant remodelling had occurred. The observed increase in succinate driven respiration in mitochondria from *nduf-11 (RNAi)* animals might be explained by one or more of the following: (1) an increased succinate entry into the mitochondria through the dicarboxylic carrier (SLC-25A10) whose expression is increased by 104%; (2) the upregulation of ubiquinone/ cytochrome c (CYC-2.1; 110%); and (3) the 25% to 75% increase in expression of Complex II subunits.

Interestingly, succinate does not fully rescue the ADP-sustained respiration when added after the Complex I-linked substrates pyruvate/ malate. Under these experimental conditions, the residual levels of Complex I in NDUF-11 depleted mitochondria are not sufficient to support maximal rates of respiration and a greater contribution from succinate is expected. However, the residual Complex I activity will lead to the production of some oxaloacetate which is an extremely potent inhibitor of Complex II [44, 45]. This will reduce rates of succinate oxidation relative to native levels and so lower the maximal respiration as observed. Although the same inhibitory effect of oxaloacetate on Complex II activity would be experienced by wild-type mitochondria, the higher levels of Complex I in this group can fully sustain maximal rates of respiration and therefore the contribution required from Complex II is minimal.

The changes observed in isolated mitochondria reflect profound physiological adaptations of the whole worm organism upon NDUF-11 depletion with a corresponding reduction of Complex I and loss of its super-complexes. Proteomic analyses indicate that affected animals also shutdown their fatty acid biosynthesis and remodel the TCA cycle towards the glyoxylate shunt. The latter is a characteristic of worm metabolism in the quiescent *dauer* state, an alternative developmental stage that promotes survival under stress, for example, when food is scarce [46]; as well as in L1 stage, prior a metabolic shift towards aerobic respiration required for entry into L2 and later developmental stages [46]. The purpose of the glyoxylate cycle is to increase nutrient stores *via* acetyl-coA conversion towards succinate through the use of fatty acids or acetate as carbon sources, which can thereafter be used in anaerobic respiration. The co-existence of impaired aerobic respiration with a shift towards anaerobic processes not only explains the downregulation of fatty acid biosynthesis, but also why the NDUF-11 depleted animals reach adulthood, albeit at slower rates.

So why do the first generation of worms with compromised Complex I organisation progress through the L2 stage instead of entering the *dauer* stage? In addition to favourable environmental cues (abundant food source), affected animals have reduced levels of DAF-18 – previously shown to be associated with defective *dauer* entry, extended lifespan and suppressed fat accumulation [47]. Moreover, *daf-18* negatively regulates germline insulin/ IGF-1 signalling (IIS) during optimal nutrient uptake [48]. Therefore, it seems plausible that the reduced mitochondrial fitness and respiratory capacity, caused by depleted NDUF-11, is sufficient for development into adulthood – albeit with reduced body size and low progeny numbers. The progeny were however sterile, most likely because the bioenergetic demands for oocyte maturation [49] and sperm motility and function [50] are such that full mitochondrial capacity is required.

Similarly, developmental arrest in the NDUF-11 knock out can be attributed to poor mitochondrial fitness resulting from loss of Complex I and its super-complexes. Maturation to the L3 and L4 stages is associated with increased energy demand, consistent with the metabolic shift from the glyoxylate shunt towards aerobic metabolism after transit through the L2 stage [51]. Moreover, knockdown of ETC subunits results in development arrest at multiple stages [52, 53]. Additionally, ubiquinone deficiency leads to L2 stage arrest [54], consistent with the importance of Complex I (and its assembled states) on worm development.

Another remarkable consequence of *nduf-11* RNAi treatment is the remodelling of the inner mitochondrial morphology. Both the qualitative and quantitative tomography data clearly demonstrate that cristae suffer from a loss of their native lamellar morphology which is likely to have detrimental effects on mitochondrial health as crista shape has been found to dictate respiratory efficiency. Significantly, aberrations in crista morphologies are associated with numerous devastating diseases including neurodegeneration and cancer [55, 56].

The perturbation of crista morphology could potentially result from reduced production of MICOS or ATP synthase components. MICOS complexes (located at crista junctions) mediate crista formation [27], and arrangement of ATP Synthase dimers into rows is instrumental in formation of sharp, curved ridges characteristic of lamellar cristae [57]. However, mass spectrometry and BN-PAGE show the latter is unaffected. In contrast, although most of the MICOS subunit levels remain constant, the one responsible for marking the nascent sites of cristae junction formation and OMM-IMM contact sites [27], Mic60 homologue IMMT-2, is significantly downregulated after reduction of NDUF-11. Interestingly also, *nduf-11* loss corresponds to widening of the cristae junctions. In spite of this there is no evident cytochrome *c* release, which explains why the bioenergetic fitness is maintained.

In conclusion, our work shows that destabilisation of Complex I and the concomitant loss of respiratory super-complex formation has a profound effect on bioenergetic capacity, mitochondrial metabolism and morphology, which severely impairs higher level physiological functions. Affected worms are unable to meet the energy requirements associated with gonad differentiation and healthy development into adulthood.

The complete ablation of NDUF-11 reveals its essential role in organismal health and proliferation, which we attribute to a catastrophic loss of Complex I activity. The milder effects of depletion, which supported *C. elegans* viability to the adult stage, allowed us to examine the consequences of Complex I destabilisation and loss of respiratory super-complexes. The prognosis for health, while not lethal, remains severe – as we have shown. We speculate that the extent of Complex I destabilisation and loss of super-complexes will have a corresponding effect on the severity of human mitochondrial disease. Indeed, patients presenting with genetic abnormalities associated with *ndufa11* present with encephalocardiomyopathy and fatal infantile lactic acidemia [17].

The effects of perturbed *ndufa-11* expression described here, provide a molecular basis for these, and potentially other afflictions involving super-complex breakdown. Perhaps also there are indirect consequences of Complex I super-complex perturbation to be considered. We show that the respiratory rebalancing for restorative oxidative phosphorylation, by increased expression and activity of Complex II, comes with the cost of excessive ROS production. It may well be that the side-effects of this physiological ‘cure’ for problems in Complex I and super-complex stability may themselves be problematic, and thus responsible for deterioration of health.

## MATERIAL AND METHODS

### Strains and nematode culture

The *C. elegans* strain N2 Bristol was used as the wild-type strain. Additional strains were obtained from the Caenorhabditis Genetics Center (CGC) (University of Minnesota, MN, US): SJ4103 (*zcIs14* [*myo-3*::GFP(mit)]); CGC43 *unc-4*(e120)/*mnC1*[*dpy-10*(e128) *unc-52*(e444) umnIs32] II. The CRISPR-Cas9 generated *cr51* knockout allele of *B0491*.*5* (this paper) was balanced *in trans* with *umnIs32*. Bacterial strains, *E. coli* strain OP50 and NA22 were used for worm propagation and *E. coli* strain HT115 was used for feeding RNAi.

*C. elegans* stocks were maintained at 20°C on NGM agar plates using standard methods [58]. Bulk liquid preparations of *C. elegans* were fed a suspension of NA22 bacteria in S-basal complete medium and incubated at 20 °C in a shaker incubator at 200 rpm. Synchronous worm populations were obtained by collecting eggs from adult hermaphrodites after alkaline hypochlorite treatment and allowing them to hatch overnight into L1 larvae in M9 buffer (3.0 g/L KH_2_PO_4_, 6.0 g/L Na_2_HPO_4_, 0.5 g/L NaCl, 1.0 g/L NH_4_Cl) lacking food [59].

### Homology modelling

Modeller 9.12 [60] was used to obtain a homology model of B0491.5 based on the NDUFA11 protein structure found in respirasome structure pdb:5GUP [19]. 5000 individual models were constructed and scored according to their Discrete Optimised Protein Energy (DOPE) value. The top 1% models were retained and analysed using the Gromacs [61] tool gmx cluster to identify clusters of well-scoring structures. The model with the best score from the largest cluster was selected.

### Molecular dynamics simulations

The stability of the homology model was validated using atomistic molecular dynamics simulation. The protein was described using the CHARMM36 force field [62], and embedded into a POPC membrane and explicit solvent using CHARMM-GUI [63, 64]. The system was energy minimized using the steepest descents method, then equilibrated with positional restraints on heavy atoms for 100 ps in the NPT ensemble at 310 K with the V-rescale thermostat and semi-isotropic Parrinello-Rahman pressure coupling [65, 66]. A production simulation was run using 2 fs time steps over 200 ns, using GROMACS 2018 [61]. Root-mean-square deviation (RMSD) and root-mean-square fluctuation (RMSF) analyses were run using the Gromacs tools gmx rms and gmx rmsf. All molecular models were represented using PyMOL.

### CRISPR-Cas9

The *cr51* knockout allele of *B0491*.*5* was obtained by CRISPR-Cas9 following the *dpy-10* co-conversion protocol developed by the Fire lab [67]. Targeted deletion of *B0491*.*5* was achieved using a pair of guide RNA (gRNA) clones pPK874 and pPK875. These were obtained by ligating complementary primer pairs (PK1801/PK1802 and PK1803/PK180; Table 2) with asymmetric Bsa I overhangs into the Bsa I site of pRB1017. The single-strand PK1805 oligodeoxynucleotide (ssODN; Table 2) repair template shares sequence homology extending beyond both the selected *B0491*.*5* PAM sites and was designed to replace the deleted region with a 7 nt sequence to terminate any potential translational read-through (Fig. S4). DNA sequencing (MWG Eurofins) confirmed the presence of a 146 bp deletion disrupting exons 1 and 2 and insertion of the 7-nt sequence.

**Table 2.**
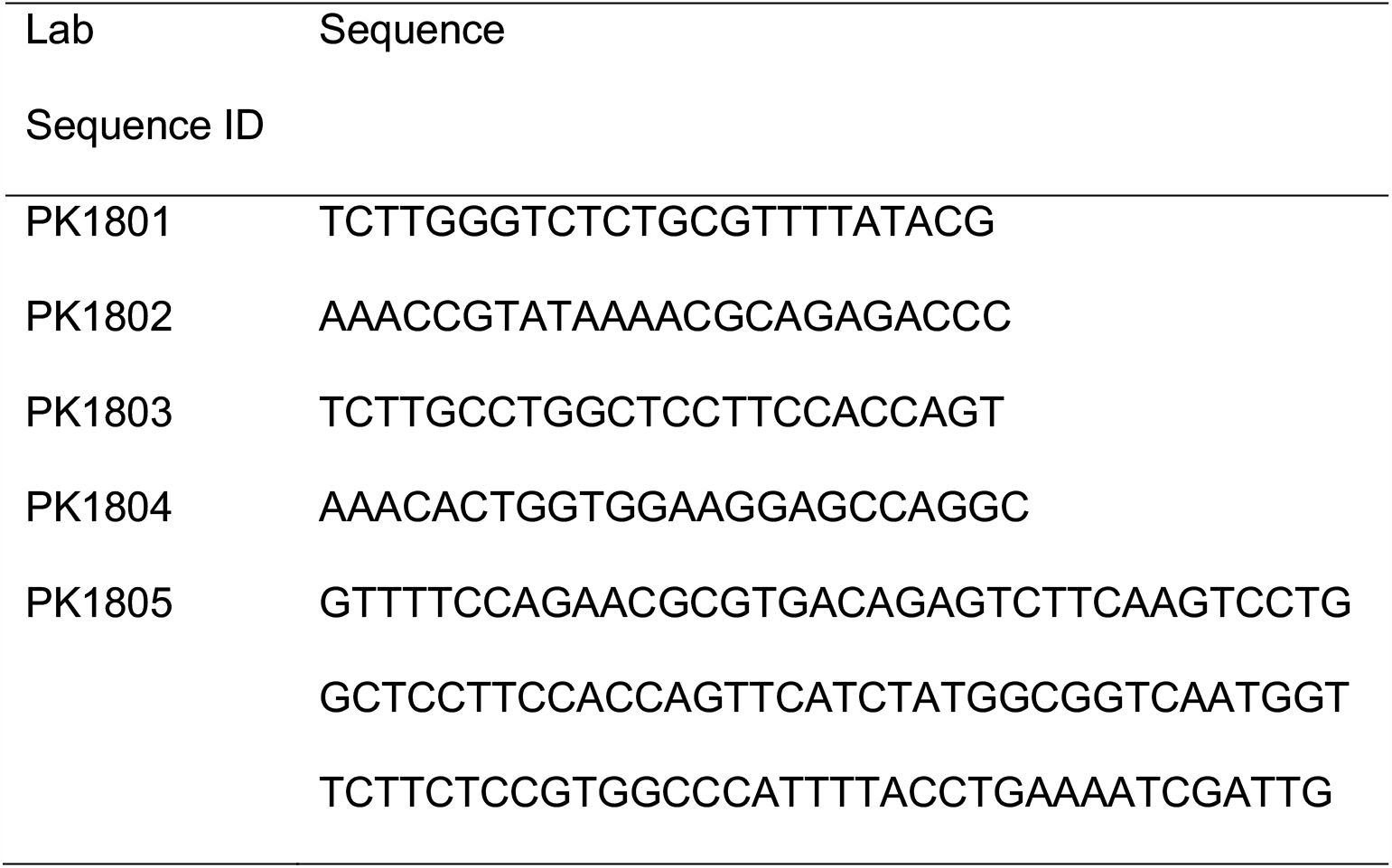
Oligonucleotides

### RNA interference

RNA interference (RNAi) was performed by feeding using clones obtained from an RNAi feeding library [68] (Source Bioscience, Nottingham, UK) and an empty vector control. To perform RNAi, synchronised *N2* L1 larvae were grown on *B0491*.*5* (*RNAi*) or empty-vector NGM feeding plates. Individual animals were plated onto fresh RNAi plates when they reached the L4 stage. Adult RNAi treated worms were then transferred every 12 h to fresh RNAi plates until egg-laying ceased. Progeny were examined and counted 24 h after the transfer of adults to fresh plates.

To obtain sufficient quantities of RNAi treated worms for biochemical analyses, 4 x 1 L cultures of 2xTY were each inoculated with 50 mL of a saturated culture of *B0491*.*5* RNAi feeding bacteria and induced with 0.4 mM IPTG after reaching an OD_600_ = 0.4. Induced cultures were grown for an additional 2 h before being harvested and resuspended in half of their original volume in S-basal complete medium supplemented with 100 μg/mL ampicillin and 0.4 mM IPTG. Cultures were inoculated with a synchronous population of L1 larvae (∼4 × 10^6^ worms) and grown in a shaker incubator at 200 rpm at 20°C for 4 days or until they reached the adult stage.

### Lifespan

Lifespan assays were performed in the presence of 5-fluorodeoxyuridine (FUdR), as described [69]. L4 stage larvae were placed on FUdR plates seeded with RNAi bacteria and incubated at 20°C. In survival curves, Day 1 represents the first day of adulthood. Animal viability was scored every other day by tactile stimulation. Animals that were lost or ruptured were not included in analyses [22]. Survival data was analysed using a Kaplan-Meier plot.

### Mitochondrial isolation

Worms were harvested from liquid culture and separated from debris by sucrose floatation [59]; ∼2-6 mL packed worm volume per 0.5 L of culture were obtained. Worms were washed extensively in M9 buffer (3.0 g/L KH_2_PO_4_, 6.0 g/L Na_2_HPO_4_, 0.5 g/L NaCl, 1.0 g/L NH_4_Cl) to restore osmolarity before being resuspended in 10 mL collagenase buffer (100 mM Tris-HCl, pH=7.4; 1 mM CaCl_2_) with 1U/mL collagenase (Sigma C0773) and incubated for 2 h at 20°C with gentle agitation. After incubation, worms were harvested by centrifugation at 160 xg and washed in STEG/M buffer (220 mM mannitol, 70 mM sucrose, 5 mM Tris-HCl, pH = 7.4, 1 mM EGTA before being resuspended in 10 mL of STEG/M(+) [STEG/M supplemented with 1 mM PMSF in methanol and 1% fatty acid-free BSA (Sigma A6003)]. Worms were homogenised in a 50 mL fitted glass/Teflon power-driven Potter-Elvejhem homogeniser before being centrifuged at 750 xg for 15 min, 4°C. The supernatant was transferred to a fresh tube and the pellet was re-extracted before pooling supernatants and collecting mitochondria by centrifugation at 12,000 xg for 15 min at 4°C. Mitochondria were resuspended and washed in 30 mL STEG/M buffer before centrifugation at 750 xg for 15 min, 4°C. The supernatant was centrifuged at 12,000 xg for 15 min, 4°C and the final pellet containing mitochondria was resuspended in 150 μL STEG/M buffer. Protein concentration was determined using a BCA protein kit (Thermo Fisher Scientific). Mitochondria were flash frozen in liquid N2 before storage at −80°C or used no later than 4-6 h post-isolation for physiological assays.

When collecting samples for mass spectrometry, worms were processed as described above and samples were harvested at specific steps during the protocol. For the cytosolic fraction, sample was collected from the supernatant after the first high-speed spin, therefore not including nuclear or membrane proteins. For the mitochondrial fraction, a sample was collected from the pellet of the first high-speed spin after performing a single wash.

### Confocal fluorescence microscopy

Worms were placed in a 5 μL drop of 1 mM levamisole on a 4% agarose pad. After 30 min, a coverslip was placed over the worms. Images were collected using a Leica TCS SP8 AOBS confocal laser scanning microscope. Images were analysed on ImageJ software (Bethesda, MD, USA).

### Transmission electron microscopy

Worms were immersed in NA22 *E. coli* media in a 0.2 mm deep gold-coated copper carrier prior to high-pressure freezing with a Leica EMpact2 high-pressure freezer (Leica Microsystems, Viena, Austria) [70]. Worms were freeze substituted for 2 h at −90 °C with 1% osmium tetroxide and 0.1% uranyl acetate in acetone using a Leica EM AFS2 freeze substitution processor (Leica Microsystems). After a temperature ramp to 0°C, specimens were washed with a graded series of acetone/Epon resin, followed twice by Epon resin alone [70] and were polymerised at 60°C between 2 Aclar (polychlorotrifluoroethylene) sheets to flatten worms and facilitate longitudinal sectioning. 1 μm thick sections were obtained and stained with Methylene Blue prior to obtaining ultrathin 70 nm sections using a Reichert Ultracut S ultramicrotome (Leica Microsystems). Sections were stained with lead and uranyl salts and images obtained using a Tecnai T12 microscope (Thermo Fisher Scientific).

### Cryo-ET of isolated mitochondria

Holey carbon EM grids (Quantifoil, Jena, DE) were glow-discharged under vacuum using an ELMO TM Glow Discharge System (Cordouan Technologies, FR) according to manufacturer’s instructions. The isolated mitochondrial suspension (5-10 mg/mL) was mixed 1:1 with a stock solution of 10 nm protein A-gold fiducial markers (Aurion, Wageningen, NL) and 3 Dl immediately applied to a glow-discharged grid. Grids were blotted for 1-1.5 sec, followed by plunge-freezing in liquid ethane using a Vitrobot Mark IV (Thermo Fisher Scientific). Samples were placed in a grid storage box and stored under liquid nitrogen prior to viewing.

Cryo-ET was performed using a FEITM TalosTM, equipped with a 200 kV FEG (Thermo Fisher Scientific), K2 DED and GIF BioQuantum LS energy filter (Gatan, Pleasanton, USA). Dose-fractionated tomograms were typically collected from +60°to 60° with tilt steps of two degrees. Gold fiducial markers were used to align tomograms and volumes were reconstructed using the IMOD software [71]. Tomogram contrast was enhanced by non-linear anisotropic diffusion (NAD) filtering in IMOD.

### Mitochondrial segmentation and morphometric analysis

Mitochondrial membranes were segmented in IMOD, through drawing and interpolation of closed contours. Using the imodmesh algorithm, contours were meshed at a z-increment of 4, with a point reduction tolerance of 2.

The mesh surface area of each membrane was computed by adding the areas of all triangles in the mesh; the volume inside the mesh of each membrane was computed by adding the volumes of tetrahedra formed between each mesh triangle and a central point within a capped mesh. Surface areas and volumes were normalised to that of the OM, before calculation of surface area: volume ratios. For display purposes, point reduction tolerance and z-increment were modified for the inner and outer membrane to give a smooth appearance, but these values were not used in calculations.

The diameter of the IMS (at 4 locations) and each segmented in-plane crista junction (CJ) was measured manually at the top, bottom and middle of each mitochondrion/junction, and an average calculated. The number of CJs in each model file was normalised to the OM surface area by calculating the frequency per nm^2^.

All data for wild-type and *nduf-11*(RNAi) tomograms were compared using unpaired, parametric Student’s t tests.

### Quantitative Mass Spectrometry

#### TMT labelling and high pH reversed-phase chromatography

100 μg aliquots of isolated mitochondria were digested with trypsin (40:1 protein:protease ratio) overnight at 37 °C and labelled using the Tandem Mass Tag (TMT) 10-plex reagent, according to the manufacturer (Thermo Fisher Scientific, Loughborough LE11 5RG, UK), and the labelled samples were pooled. Sample handling and fractionation by high pH reversed-phase chromatography was carried as previously described [72].

#### Nano-LC mass spectrometry

High pH RP fractions were further fractionated using an Ultimate 3000 nano-LC system in line with an Orbitrap Fusion Tribrid Mass Spectrometer (Thermo Scientific), as previously described [72].

Raw data files were processed and quantified using Proteome Discoverer software v2.1 (Thermo Scientific) and searched against the UniProt *Caenorhabditis elegans* database (downloaded in December 2018: 27626 entries) using the SEQUEST algorithm. Peptide precursor mass tolerance was set at 10 ppm, and MS/MS tolerance was set at 0.6 Da. Search criteria included oxidation of methionine (+15.9949) as a variable modification and carbamidomethylation of cysteine (+57.0214) and the addition of the TMT mass tag (+229.163) to peptide N-termini and lysine as fixed modifications. Searches were performed with full tryptic digestion and a maximum of 2 missed cleavages were allowed. The reverse database search option was enabled and all data was filtered to satisfy false discovery rate (FDR) of 5%.

Statistical analysis for the quantitative mass spectrometry was carried out first by log_2_-transformind the data (protein abundance) so that the data followed near normal distribution and skewness was reduced. The log_2_-fold change was then calculated between B0491.5(RNAi) and the control samples by subtracting the mean of the former from that of the latter. Unpaired, two-tailed Students’ t-tests were performed on the transformed data to calculate the p-values. The log_2_-fold change and log_10_ of the p-values were then plotted on the volcano plots to show the distribution of the data.

### O_2_ consumption

Oxygen consumption was measured by high-resolution respirometry at 25 °C in a KCl-based respiration buffer at pH=7.3 (125 mM KCl, 10 mM Tris, 20 mM MOPS, 2.5 mM KH_2_PO_4_, 2.5 mM MgCl_2_) using a 2 mL chamber of an Oxygraph 2k (Oroboros Instruments, Innsbruck, AT). 0.25 mg/mL mitochondria in respiration buffer were added to each chamber and equilibrated before the addition of various respiratory substrates (final concentrations indicated): 10 mM L-malate/5 mM pyruvate (Complex I-linked respiration) or 5 mM succinate plus 1 μM rotenone (Complex II-linked respiration). After a stable respiration state 2 was achieved, ADP was added to 1mM (excess), followed by cytochrome *c* (10 µM). 1 μM oligomycin was used to inhibit ATP synthase and induce a pseudo-state 4. These parameters were based in part on previously published respiration experiments in *C. elegans* [73]. Respiration rates were calculated as the average value over a 30 s window in DatLab 5 (Oroboros Instruments) and expressed in nmol O_2_ / min /mg protein. Respiration states were identified according to Chance and Williams [74]. Statistical significance between groups was determined using a paired two-tailed Students’ t-test to control for day-to-day variability associated with the isolation protocol and electrode calibration.

### Mitochondrial Membrane Potential

Membrane potential (ΔΨ) assessment was carried out with the fluorescent potentiometric dye tetramethylrhodamine methyl ester (TMRM; Sigma-Aldrich) working in quench mode (1 µM). Mitochondrial depolarisation causes an increase in fluorescence. Fluorescence was recorded at room temperature with a Varian Cary 50 UV-Vis Spectrophotometer (Agilent Technologies, Santa Clara, CA, US) using 2 mL of KCl respiration medium (above) supplemented with either 1 µM TMRM and 10 mM L-malate/5 mM pyruvate (Complex I-linked respiration) or 5 mM succinate plus 1 μM rotenone (Complex II-linked respiration). After obtaining a stable signal in the absence of the biological sample, reaction was started by addition of mitochondria (0.25 mg/mL) and followed by substrates and/or inhibitors added sequentially: coupled respiration was stimulated with the addition of 1 mM ADP (state 3 respiration); ATP synthase was inhibited with 1 μM oligomycin; and, maximum depolarisation was achieved using 1 μM CCCP and 1 μM antimycin A.

No attempt was made to calibrate the fluorescent TMRM signal and thus, relative fluorescent units rather than milli-volts are shown in this work. The averaged signal of each plateau was calculated over 15 seconds using DatLab 2 in MS-DOS. Statistical significance between groups was determined using a paired two-tailed Students’ t-test to control for day-to-day variability associated with the isolation protocol.

### Hydrogen peroxide production

ROS production was assayed with the hydrogen peroxide sensitive fluorescent dye Amplex Red (Thermo Fisher Scientific), as described [75], and measured using a Varian Cary 50 UV-Vis Spectrophotometer (Agilent Technologies, Santa Clara, California, United States) with manual mixing between additions. Mitochondria at a concentration of 0.25 mg/mL were added to a 2 mL cuvette in KCl-based respiration medium. Mitochondria were energised with either 10 mM L-malate/5 mM pyruvate (Complex I-linked respiration) or 5 mM succinate (Complex II-linked respiration). Substrates were sequentially added as follows: 1 μM rotenone for inhibition at Complex I level; and, 1 μM antimycin A for inhibition at Complex III level. The averaged signal of each plateau was calculated over 15 seconds using DatLab 2 in MS-DOS. Statistical significance between groups was determined using a paired two-tailed Students’ t-test to control for day-to-day variability associated with the isolation protocol.

### LDS-Polyacrylamide Gel Electrophoresis (PAGE)

Pre-cast Bolt gels and buffers (Thermo Fisher Scientific) were used. Whole worm protein samples consisted of ∼50 adult worms in 15 μL of M9 buffer mixed with an equal volume of 4X Bolt LDS (Lithium Dodecyl Sulfate) NuPAGE sample buffer containing 250 mM 1,4-dithiothreitol (DTT). Mitochondrial samples contained 50 μg of protein diluted 1:4 with 4x Bolt LDS NuPAGE sample buffer containing 250 mM DTT. All samples were incubated at 60 °C for 25 min and centrifuged for 1 min at 14,500 xg before electrophoresis on 4-12% Bolt Bis-Tris Plus gels at 150 V (constant) in Bolt MOPS running buffer for ∼30 min. Gels were transferred to 0.45 μm PVDF membranes (Millipore) at a 25 V (constant) for 10 min using a Pierce semi-dry G2 Fast Blotter (Thermo Fisher Scientific) according to manufacturer.

### Western Blot

Membranes were incubated in BLOTTO (5% milk in TBS-T, 50 mM Tris-HCl, pH 7.6, 150 mM NaCl, 0.05% Tween-20) for ∼1 h at room temperature. Primary antibodies were diluted in BLOTTO and incubated for 1 h at room temperature with constant shaking. Membranes were washed with TBS-T for 10 min before incubating for 1 h at room temperature with a secondary antibody conjugated to goat anti-mouse HRP (NB7539; Novus Biologicals, Centennial, CO, USA) with a titre of 1:10,000 in BLOTTO. Membranes were washed three times with TBS-T for 10 min each before incubation with Amersham ECL detection reagent (GE Healthcare) and chemiluminescent detection using the Li-COR Odyssey FC imaging system (LI-COR Biosciences). Alternatively, secondary antibodies conjugated with fluorescent dyes were directly visualised at 700 or 800 nm (anti-mouse 800, Invitrogen, SA5-10172; anti-rabbit 680, ThermoFisher, 35569). Western blot quantification was performed using LI-COR image studio. Titres and antibodies used in this study include: (1:1,000) mouse anti-NDUFS3 [17D95] mAb (Abcam, ab14711) and (1:1,000) mouse anti-ATP5A [15H4C4] mAb (Abcam, ab14748). A rabbit polyclonal antibody recognising a N-terminal peptide sequence of CEL-B0491.5 (GHGEEPLTATYKTQR-[C]-amide) was produced by Cambridge Research Biochemicals (Billingham, UK) and used at a titre of 1:500.

Statistical significance between groups was determined using an unpaired two-tailed Students’ t-test.

### Blue-Native PAGE

Previously prepared mitochondria (1 or 2 mg) were thawed and pelleted at 14,500 xg for 15 min at 4 °C and resuspended in solubilisation buffer (50 mM NaCl, 10% glycerol, 1 mM PMSF and 20 mM Tris pH 7.5) supplemented with either 10 mg/mL digitonin (Millipore-Sigma, cat # 300410); or 5 mg/mL Triton X-100 (Merck, X100). Solubilisation was carried at 4 °C for 20 min with mild agitation, followed by centrifugation at 14,500 xg for 15 min at 4 °C. Solubilised proteins in a 25 μL volume were prepared and separated by electrophoresis on NativePAGE 3-12% Bis-Tris gels (Thermo Fisher Scientific) and NuPAGE Tris-Acetate buffer (Thermo Fisher Scientific) according to the manufacturer at 30 V (constant) for ∼16 hours or until the dye front reached the gel bottom. Gels were stained with either Coomassie G-250 (50% methanol, 10% acetic acid and 0.5% Coomassie G-250) and de-stained (10% methanol; 10% acetic acid) or silver stained using the Silver Quest kit (Thermo Fisher Scientific).

### In-gel respiratory complex

In-gel identification of respiratory complexes separated by BN-PAGE was performed using previously published methods [76, 77]. Briefly, BN-PAGE gels were incubated at room temperature on an orbital shaker at 60 rpm with Complex I activity buffer (50 mM potassium phosphate buffer pH= 7.4, 0.1 mg/mL NADH, 0.2 mg/mL nitro blue tetrazolium chloride). Incubation was carried out for ∼1 h.

### 2-dimensional BN/LDS-PAGE

A lane from a BN-PAGE gel was trimmed to fit the loading area of a 2^nd^ dimension LDS-PAGE gel and incubated in 10 mL of reducing solution (1x Bolt LDS NuPAGE sample buffer with 250 mM DTT) for 30 min with agitation. The gel lane was then inserted into the well of a pre-cast 4-12% NuPage Bis-Tris 1-well gel. The well was overlaid with 1x Bolt LDS buffer and electrophoresed at 150 V (constant) for 40 min or until the dye front reached the gel bottom. The gel was either silver stained (above) to visualise total protein or used for western blot analysis to detect specific proteins.

### Statistical Analysis

Data were assessed for normality using the Kolmogorov-Smirnov test with Dallal-Wilkinson-Lillie for correction. Overall, analyzed data were normally distributed. Since group sample sizes are equal and the parametric statistical tests applied are robust for moderate deviations from homoscedasticity, parametric tests were always applied [78].

The statistical tests used in this study are described in each appropriate section. Statistical analyses were performed using Graph Pad Prism version 7.0 and/or 8.0 (GraphPad Software, Inc., San Diego, CA, USA). Volcano plots were made in VolcanoR [79].

## AUTHOR CONTRIBUTIONS

Conceptualization – IC, PK, AKW;

Data Curation – AKW, GCP, EB, RAC, VAMG, CN, PK;

Formal Analysis – AKW, EB, CN, PK;

Funding Acquisition – IC;

Investigation – AKW, GCP, EB, PK;

Project Administration – IC, PK;

Resources – PV, APH, VAMG, PK;

Supervision – VAMG, PK, IC;

Writing – Original Draft Preparation – AKW, GCP, PK, IC.

## ACKNOWLEDGMENTS

This research was funded in part, by the Wellcome Trust: Investigator Award to IC (104632/Z/14/Z) and a Multi-User Equipment Grant for electron cryo-microscopy instrumentation to IC (202904/Z/16/Z) and to VAMG (206181/Z/17/Z). For the purpose of Open Access, the author has applied a CC BY public copyright licence to any Author Accepted Manuscript version arising from this submission. AKW is recipient of a University of Bristol PhD scholarship, and EB of a PhD studentship from the BBSRC SWBioDTP DTP (BB/M009122/1). We would like to thank Margherita Protasoni for useful discussions on super-complexes and their differences between species.

## Abbreviations

BN-PAGE: blue native - polyacrylamide gel electrophoresis
CL: cardiolipin
cryo-EM: cryo-electron microscopy
cryo-ET: cryo-electron tomography
ETC: electron transport chain
IMM: inner mitochondrial membrane
TMH: transmembrane helix

## SUPPLEMENTARY FIGURE / MOVIE LEGENDS

**Supplementary Figure S1.**
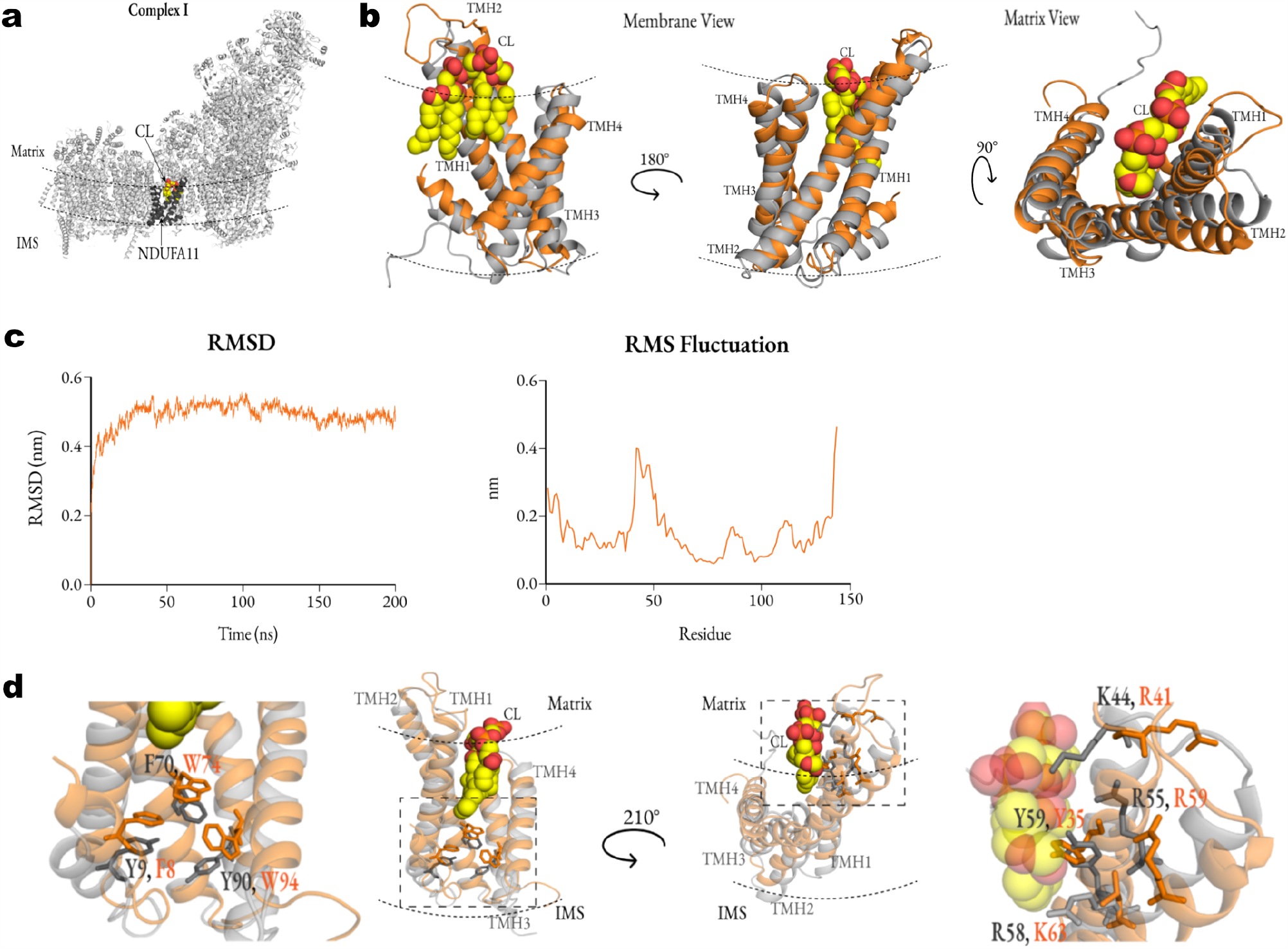
**(a)** Human NDUFA11 subunit (darker grey) on Complex I (light grey; porcine, pdb: 5GUP, [19]) used for the homology model and derived homology models of *C. elegans* NDUF-11 **(b** and **d). (b)** Three orientations of the *C. elegans* NDUF-11 model (orange) overlaid with NDUFA11 structure from 5GUP (grey). **(c)** Left: RMSD plot of the *C. elegans* NDUF-11 protein backbone over 200 ns of atomistic simulation. The protein is very stable, considering that the input is a homology model. Right: RMS fluctuation of each residue over the simulation, fitting to the input structure. The main protein is very stable, with a flexible loop between TMH1 and TMH2, and a flexible C-terminus. **(d)** Cardiolipin interacting residues shown in membrane view of different orientations; zoomed regions of the dotted box are shown on the side of the central panels. Residues labelled in the figure correspond to the NDUFA11 structure (grey) and NDUF-11 homology model (orange). All structures were created in PyMOL and assembled in a photo editor; the dotted lines represent the membrane border. Abbreviations: CI – Complex I, CIII_2_ – dimer of Complex II, CIV – Complex IV, CL – cardiolipin, IMS – intermembrane space, THM – transmembrane helix.

**Supplementary Figure S2.**
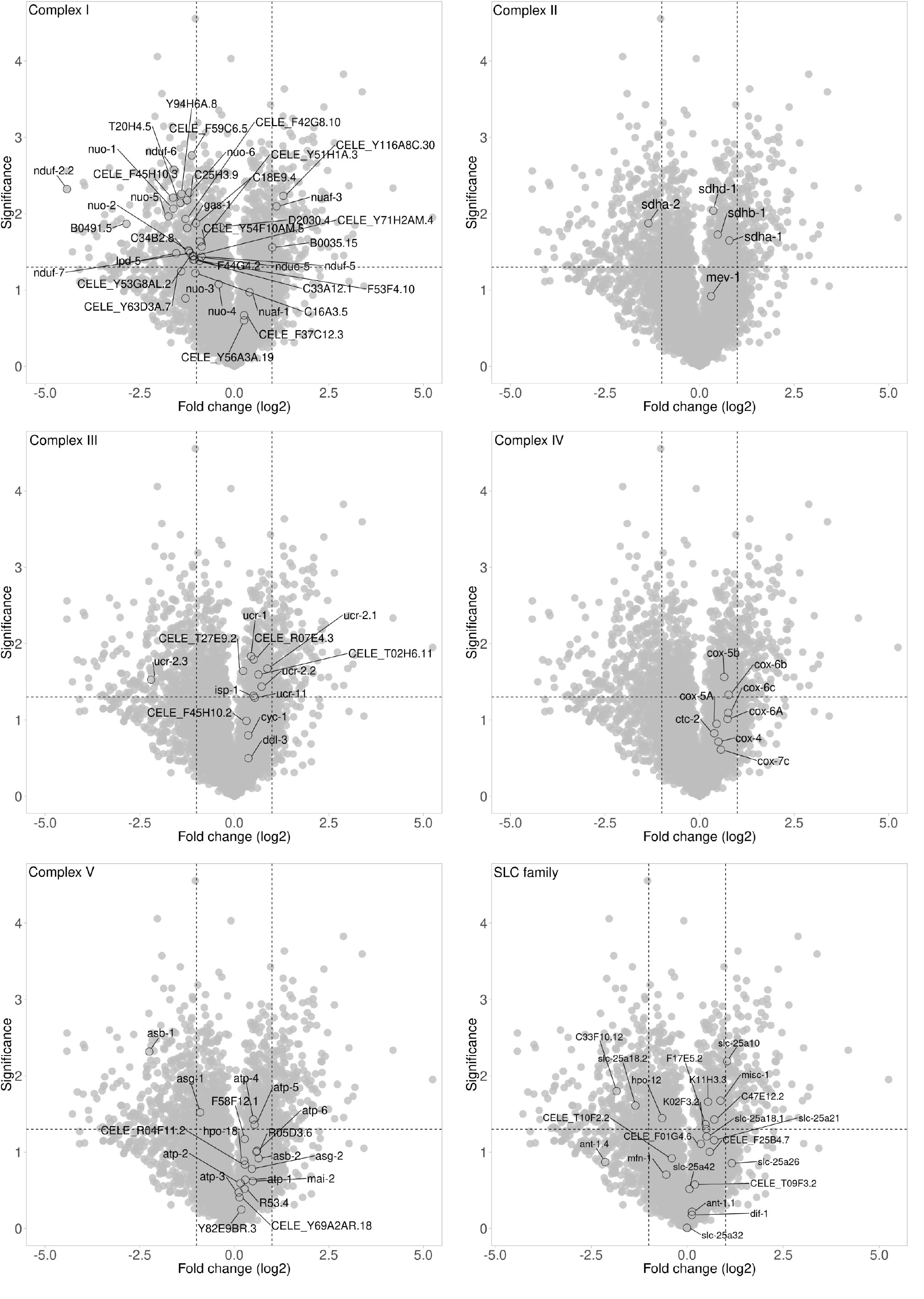
Volcano plots showing mass spec data from isolated mitochondrial fractions as shown in Fig. 3c. In each panel, labelled data represent subunits of OXPHOS complexes as shown on the top left corner of each graph; and, members of the solute carrier superfamily (SLC). Vertical dashed lines represent the user-defined threshold in expression levels while horizontal dash lines represent the p-value of 0.05. Therefore, the most upregulated genes are towards the right, the most downregulated genes are towards the left, and the most statistically significant genes are towards the top. Plots are in log_2_-log_2_ scale and were generated using VolcanoR.

**Supplementary Figure S3.**
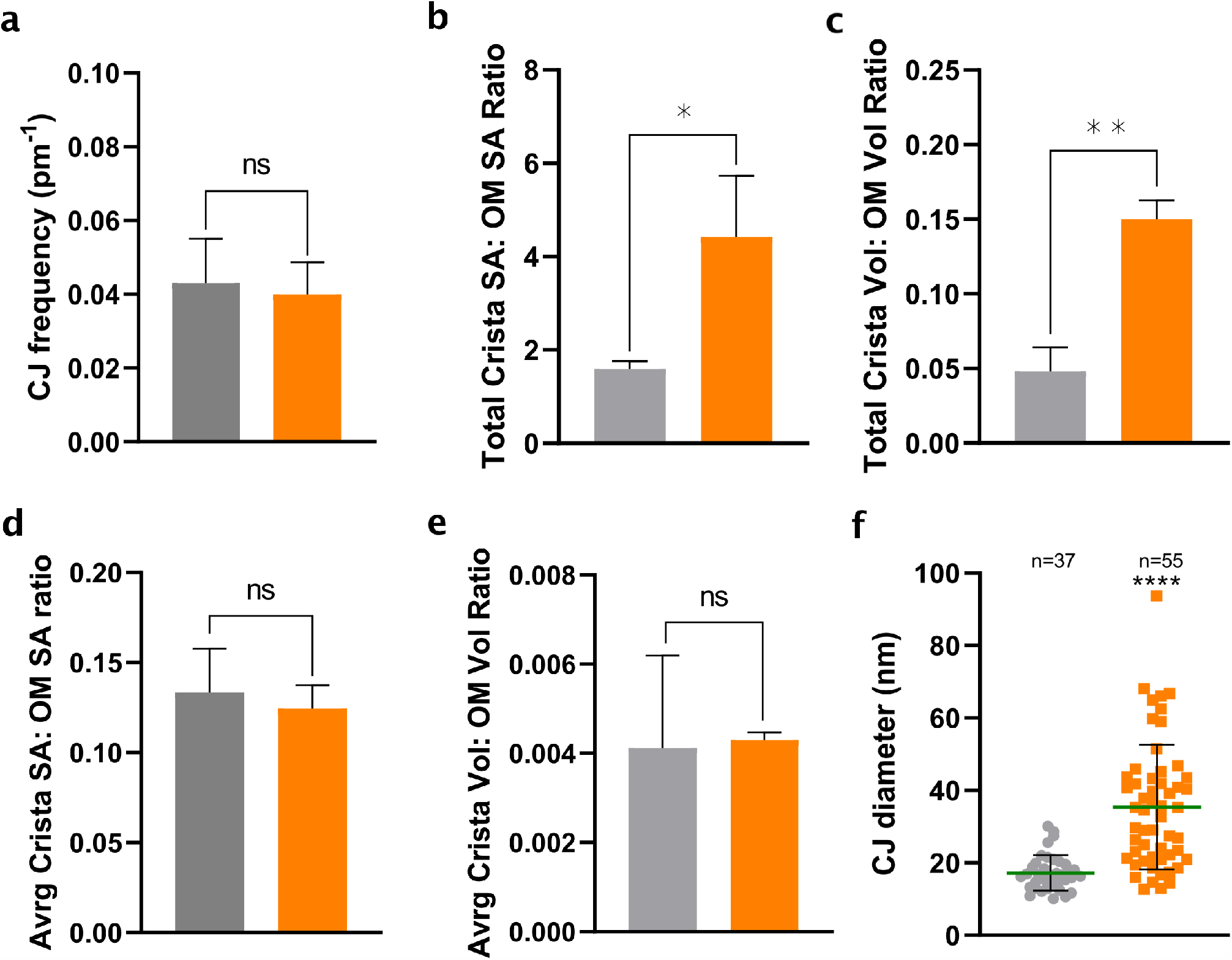
Effect of NDUF-11 knockdown on the total and average crista surface areas and volumes, and CJ frequency and diameter. (**a**) Effect of *nduf-11* knockdown on CJ frequency. The number of CJs in the segmented models from main Fig. 6a-f were counted, and normalised to the outer membrane surface area. **(b**,**c)** Total crista surface area and volume presented as a ratio to outer membrane surface area and volume, respectively. There is an increase in the total surface area and volumes of crista membranes relative to mitochondrial size for the NDUF-11 knockdown. **(d**,**e)** Average crista surface area and volume presented as a ratio to the outer membrane surface area and volume, respectively. Mesh surface area and volume inside the mesh of each membrane from mitochondrial reconstructions shown in main Fig. 6a-h were calculated computationally (n=3 for each condition, and n = 37 (wild-type) and 105 (NDUF-11) for total crista analysed). There is no significant change in the average surface area and volume per crista membrane relative to mitochondrial size, representative of a mix of fused and fragmented cristae in the NDUF-11 knockdown. **(f)** Effect of NDUF-11 knockdown on distribution of CJ diameters. CJ diameter was measured from mitochondrial reconstructions. The diameters of 37 junctions from three wild-type mitochondria shown in main Fig. 6a-c, and 55 junctions from three nduf-11(RNAi) mitochondria shown in main Fig. 6d-f were plotted individually. Unpaired, parametric t-tests were used to calculate all significance values: * p ≤ 0.05, ** p ≤ 0.01. Error bars: SEM in panel b-c, SD in others.

**Supplementary Figure S4.**
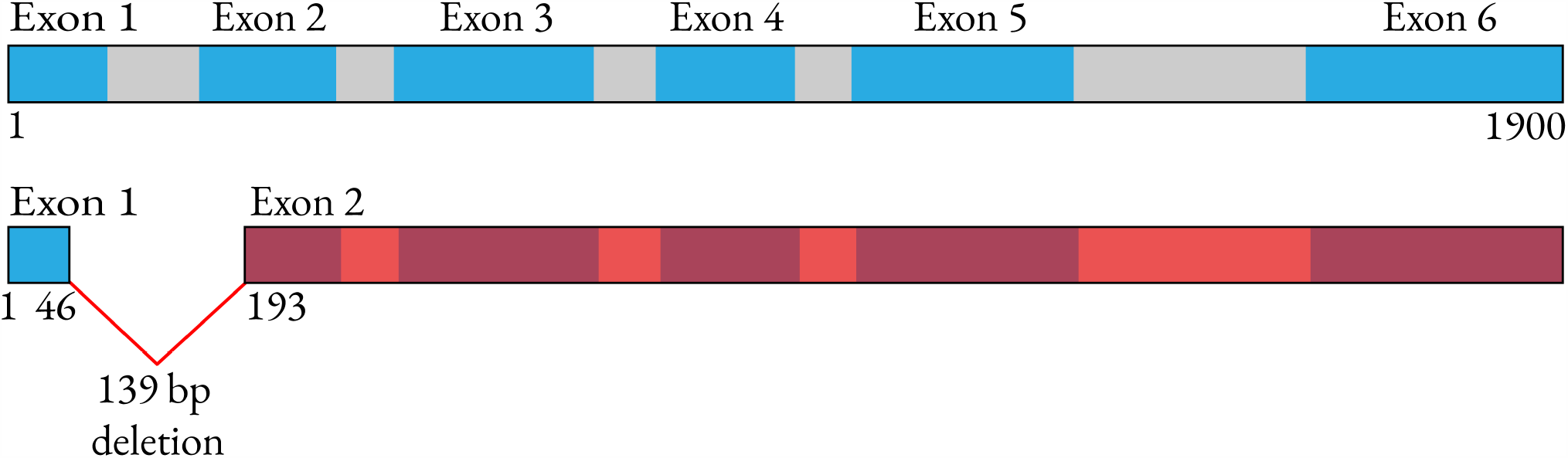
*nduf-11*(*cr51*) CRISPR-Cas9 mutant - schematic representation of the CRISPR mutation introduced to *nduf-11*, leading to a deleted region in between position 46 and 193 and thus a frameshift mutation which perturbs the rest of the sequence represented in red.

**Supplementary Movie M1 and M2 –** Movies showing a 360° rotation about the y-axis of segmentations from Fig. 6a (movie M1; control sample) and Fig. 6f (movie M2; RNAi sample), respectively. The outer membrane is hidden to display crista junction morphology. An image sequence of 100 PGN files was collected in IMOD, and sequence montaged into 10fps mp4 file in ImageJ.

